# Fragment screening using biolayer interferometry reveals ligands targeting the SHP-motif binding site of the AAA+ ATPase p97

**DOI:** 10.1101/2022.06.01.494394

**Authors:** Sebastian Bothe, Petra Hänzelmann, Stephan Böhler, Josef Kehrein, Christoph Wiedemann, Ute A. Hellmich, Ruth Brenk, Hermann Schindelin, Christoph Sotriffer

## Abstract

Biosensor techniques have become increasingly important for fragment-based drug discovery during the last years. Here, we describe a biolayer interferometry-based fragment screen targeting the AAA+ ATPase p97, an essential protein with key roles in protein homeostasis and a possible target for cancer chemotherapy. Currently available p97 inhibitors target its ATPase activity and globally impair p97-mediated processes. In contrast, inhibition of cofactor binding to the N-domain by a protein-protein-interaction inhibitor would enable the selective targeting of specific p97 functions. We demonstrate that a region known as SHP-motif binding site can be targeted with small molecules. Guided by molecular dynamics simulations, the binding sites of selected screening hits were postulated and experimentally validated using protein- and ligand-based NMR techniques, as well as X-ray crystallography, ultimately resulting in the first structure of a small molecule in complex with the N-domain of p97. The identified fragments provide insights into how this region could be targeted and present first chemical starting points for the development of a protein-protein interaction inhibitor preventing the binding of selected cofactors to p97.

## Introduction

The AAA+ ATPase p97 is an essential protein involved in numerous cellular processes and plays multiple key roles during protein homeostasis [1,2]. p97 consists of six identical monomers that assemble into a functional C_6_-symmetrical hexamer. Each monomer can be subdivided into an N-terminal domain, the Walker A and Walker B motif-containing ATPase domains D1 and D2, and an extended unstructured C-terminal tail (**Fig. 1a**). The ATPase functions convert the energy of ATP hydrolysis into mechanical energy required to extract ubiquitinylated proteins from membranes, chromatin or macromolecular complexes [1]. The functional diversity of p97 is mediated through the interaction with a large number of distinct protein cofactors, which mainly interact with the N-terminal domain of p97 and also the C-terminal tail [3,4]. Cofactors binding to the N-domain interact with two different regions. UBX (ubiquitin regulatory X) or UBXL (UBX-like) domain containing cofactors as well as cofactors harboring a VIM (VCP-interacting motif) or VBM (VCP-binding motif) bind in a cleft between the Nn- and Nc-subdomains of the N domain. In contrast, cofactors with a SHP-motif interact with an extended region located in the Nc-subdomain [5].

**Figure 1.**
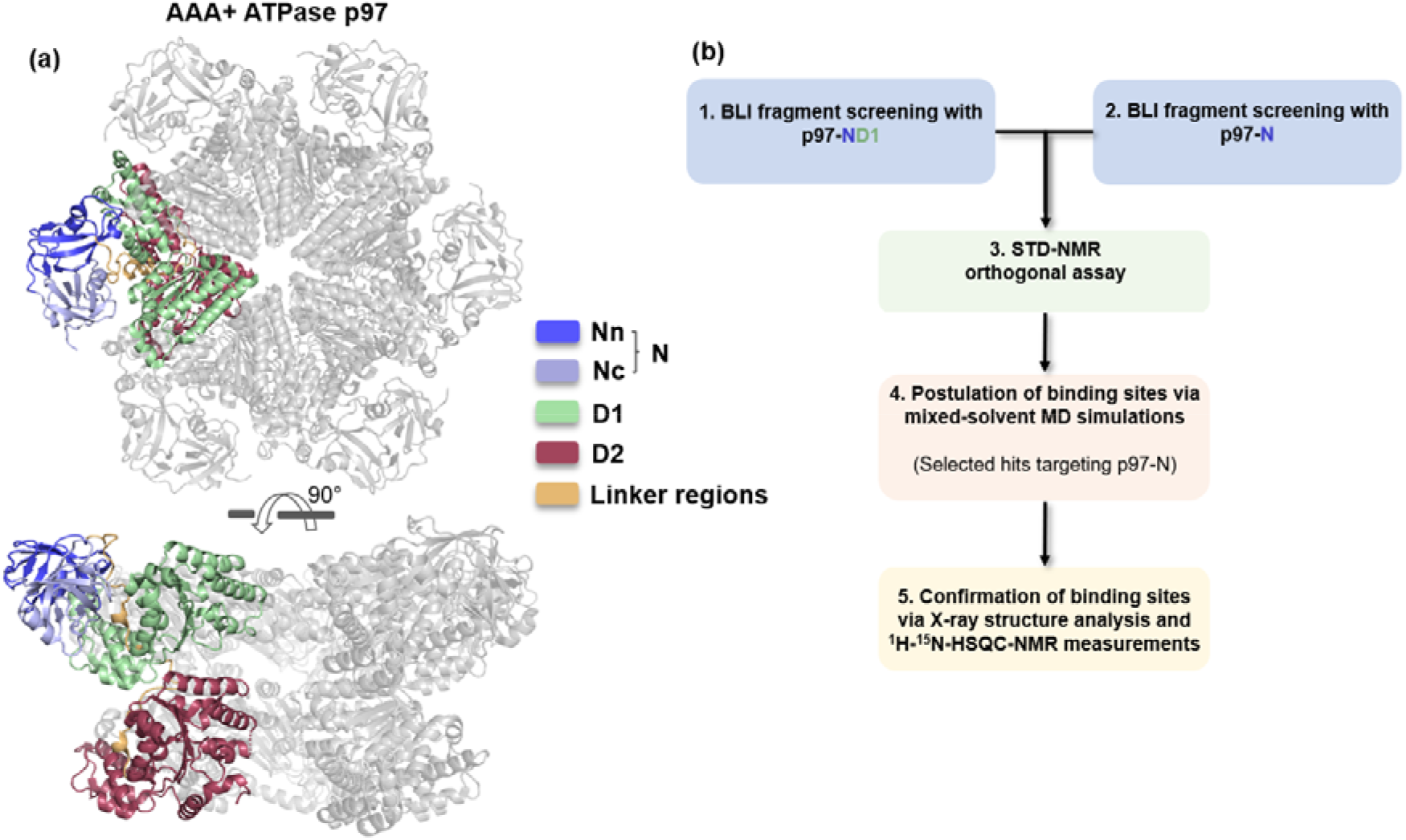
Structure of p97 and workflow of the fragment screen. **(a)** Hexameric assembly of p97 (PDB: 5C1B) with one monomer shown in color. Each monomer can be divided into the two ATPase domains, D1 (green) and D2 (red), and an N-terminal domain consisting of two subdomains, Nn (blue) and Nc (light blue), where most of the known cofactors bind. The domains are connected by two linker regions (orange). **(b)** Workflow of the fragment screen. To validate the screen, it was first performed using a p97-ND1 construct. The established experimental conditions were transferred to the isolated N-domain, followed up by an orthogonal STD-NMR assay and the identification of the binding region of selected hits.

Point mutations in p97 lead to IBMPFD (inclusion body myopathy associated with paget’s disease of the bone and frontotemporal dementia) [6] and FALS (familial amyotrophic lateral sclerosis) [7]. Furthermore, due to its significant role in regulating a variety of physiological responses, especially as part of the ubiquitin-proteasome-system (UPS), p97 has emerged as a potential therapeutic target, in particular to treat cancer [8]. Inhibition of p97 leads to cell death in HCT116, A549 [9] and B-ALL [10] tumor cells.

Based on their mechanism of action, known p97 inhibitors can be divided into three categories [7,11,12]: (i) Competitive D1 and/or D2 ATPase inhibitors (e.g., DBeQ [13], ML240/241 [14], NMS-859 [15] or CB-5083 [16]); (ii) allosteric inhibitors of the ATPase function by a non-competitive mechanism (e.g. NMS 873 [17], UPCDC30245 [18] or MSC1094308 [19]); and (iii) inhibitors with unknown mechanism of action (e.g. clotrimazol [20] or eeyarestatin I [21]). Furthermore, inhibitors can be divided into non-covalent ligands and covalent binders, the latter mainly modifying residue C522 in the nucleotide binding site of the D2 domain [11,15]. So far, only the natural product xanthohumol, occurring in hop plants, has been reported to bind the N-domain of p97 [22].

As demonstrated by the failure of the CB-5083 inhibitor in clinical phase I trials [12], inhibitors that competitively target the ATPase functions of p97 may exhibit limited selectivity due to the inhibition of other nucleotide-dependent enzymes. Furthermore, inhibition of the ATPase function leads to an indiscriminate loss of cellular p97 functions. In contrast, inhibition of cofactor binding to the N-domain by a protein-protein-interaction (PPI) inhibitor would enable the selective targeting of specific p97 functions. Beyond the long-term goal of using them as therapeutics, these inhibitors would also be important to dissect the molecular and cellular functions of p97 and its cofactors, thus helping to unravel how certain cofactors control p97 activity.

To identify chemical starting points for the development of PPI inhibitors targeting the N-domain of p97 (p97-N), we used a fragment-based screening approach (**Fig. 1b**). In contrast to screenings with drug-like compounds, the use of “fragments” (i.e., smaller compounds with 10–20 heavy atoms) can sample the regions of chemical space more efficiently [23]. A fragment-based approach is also best suited to address more challenging targets, such as interfaces of protein-protein complexes, and has already been used in the successful development of PPI inhibitors against IL-2 [24], TNF-α or BCL-X_L_ [25]. In the single hitherto reported fragment screening with p97 [26], six fragments binding to the N-domain were identified by surface plasmon resonance spectroscopy (SPR) and saturation transfer difference (STD)-NMR methods.

The screen reported here was conducted by biolayer-interferometry (BLI), followed up by STD-NMR experiments. To map the binding sites of the identified fragments we used mixed-solvent molecular dynamics (MD) simulations as well as X-ray crystallography and ^1^H-^15^N-HSQC (heteronuclear single quantum coherence) NMR measurements to confirm the binding sites predicted by the simulations. Importantly, the first structure of a small molecule in complex with the N-domain of p97 could be obtained by X-ray crystallography, presenting a starting point for further optimization and structure-based drug design. Moreover, this study demonstrates that the BLI technique, which has been reported so far only sporadically in the literature [27], is well suited for detecting fragments and is a viable option besides the commonly used SPR method.

## Results

### Initial biolayer interferometry screenings

As starting point for the initial screen, a BLI binding assay was developed with a p97-ND1 construct (amino acids 2-481) and validated with ADP as a positive control. The p97-ND1 construct was enzymatically biotinylated via an Avi-Tag and *E. coli* biotin ligase [28] and loaded to high response (>5 nm) on Super Streptavidin Biosensors (SSA), yielding a homogeneous sensor surface. The affinity (K_d_) of ADP to p97-ND1 was measured to be 200 ± 14 nM using a Langmuir model based on steady state measurements of a dilution series. This value is in agreement with reported affinities for ND1 constructs of similar lengths; e.g., a value of 200 nM was reported for an ND1 fragment spanning amino acids 1-458, and a value of 98 nM for an ND1 fragment spanning amino acids 1-480 [26,29]. The kinetic analysis of the sensorgrams indicated a clear 1:1 binding model for ADP. The global exponential regression resulted in an affinity value of 239 ± 9 nM, in agreement with the previous results. A comparison of the kinetic rate constants determined by SPR measurements with a comparable ND1 construct [29] revealed that the *k_on_* and *k_off_* values measured for ADP by BLI are in a similar range (see **Fig. S1**). The surprisingly high responses upon ADP-binding with shifts of more than 0.7 nm indicate that the BLI signal is not only dependent on the molar weight (as in SPR), but may also reflect conformational changes induced in p97 upon ADP binding.

For hit discovery, the BiSS fragment library containing 679 fragments was screened (cf. Materials and Methods). Due to the lack of a positive control for p97-N, the fragment library was first screened with p97-ND1 to validate the experimental conditions. The screen resulted in an overall Z-value [30] of 0.67, demonstrating its high quality (values between 0.5 and 1.0 represent excellent assays). A fraction of 82 fragments was identified above the chosen threshold (> 1.0 sigma, **Fig. S2**). Fragments showing abnormally high or negative signals or untypical shapes of the BLI curves were eliminated after visual inspection, leaving 48 (6.97%) compounds for subsequent hit validation. Out of these, 29 fragments exhibited a dose-dependent response, demonstrating the functionality of the assay. Therefore, the same conditions were transferred to a screen with p97-N and the aforementioned library. 66 fragments were above the threshold (> 1.0 sigma), and 42 (6.52%) compounds were selected for dose-response experiments after visual inspection of the sensorgrams (**Fig. S2**).

### Biolayer interferometry dose-response analysis of N-domain hits

The selected 42 fragments of the screen with the N-domain were tested in a dose-dependent manner using concentrations from 1000.0 µM to 31.3 µM in 1:1 dilution steps. 36 compounds exhibited dose-dependent response signals. While some sensorgrams displayed behavior characteristic of unspecific binding, 22 curves displayed binding curves indicative of fragment binding (**Fig. S3**). The sensorgrams of these fragments were fitted with a 1:1 Langmuir model to estimate their affinities and analyzed with respect to the following parameters: (i) The presence of a stationary phase; (ii) curve shapes according to either a homogeneous (1:1) or a heterogeneous (2:1) binding model, and (iii) justification of a 1:1 binding model via the obtained empirical constants *k_obs_* of the association phase. Based on these analyses, a Score_BLI_ was calculated (for details see Supplementary Methods in the Supporting Information). The calculated Score_BLI_ should be close to or above 1.0 for ligands that can be described with a 1:1 model, whereas ligands showing unspecific binding yield values << 1.0. Based on the Score_BLI_, we divided the fragments into a low-scoring (< 0.70) and a high-scoring (≥ 0.70) group.

### Confirmation of selected fragments by STD-NMR

To confirm the hits of the BLI screening, fragments with p97-N K_D_ values below 700 µM were investigated by STD-NMR. A total of 19 fragments satisfied this affinity cutoff. The STD-NMR method provides insights into the transient binding of a small molecule to a protein in solution. Therefore, the method is suitable to eliminate false positive hits arising from the immobilization of the target protein in the BLI-based screen. The NMR measurements were carried out with the biologically more relevant p97-ND1 construct in the presence of 500 µM ADP to block the nucleotide binding site. The STD effect was calculated by dividing the intensity of the difference spectrum by the intensity of the off-spectrum (I_diff_/ I_off_ x 100) for each signal [31]. For further analysis only resonances in the aromatic region of the ^1^H NMR spectrum of the individual ligand were considered. A *mean STD* effect of each fragment interacting with p97-ND1 was calculated using the sum of the individual STD-effects divided by the number of signals considered. Furthermore, the *maximum STD*-effect of each compound was determined. Three out of the 19 fragments exhibited no significant effect (S/N < 3.0) and were therefore not confirmed by the STD-NMR assay. The remaining fragments showed *maximum* STD-effects in a range from 10.5% (TROLL9) up to 72.5% (TROLL6). Similar results were obtained in the previously reported fragment screen with p97 [26]. An overview of the STD-effects is shown in **Fig. S4.**

### Selection of the identified fragments for further analysis

For further selection of the fragments the Score_BLI_ was used as criterion. All fragments showing a Score_BLI_ value greater than 0.7 were further investigated, resulting in 13 molecules (**Fig. S4**). To rule out the possibility that the fragments bind to regions of the N-domain that are normally blocked by the D1-domain of p97, dose-dependent measurements were carried out with the ND1 construct, again using the BLI method. For 12 fragments, dose-dependent binding was observed and a 1:1 Langmuir model could be applied to obtain the respective K_D_ value. The determined K_D_^(p97-ND1)^ values mostly agreed well with those previously determined for the respective fragments with the isolated N-domain. Only TROLL1 showed no response with the p97 ND1-construct, and VIK40 bound the ND1-construct with only low affinity (K_D_ > 700 µM). TROLL1 and VIK40 were therefore not considered further.

The sensorgrams of the fragments showed surprisingly slow on and off rates. Therefore, based on the kinetic data the affinity was determined and plotted against the affinity deduced from the steady state analysis (**Fig. S5**). 7 fragments showed comparable affinities indicating that the observed slow on and off rates result from the BLI assay. 3 fragments (TROLL13, TROLL14 and TROLL18) showed a higher discrepancy for the calculated affinities with up to 3.5 times lower affinity deduced from the kinetic data. TROLL10 showed the highest discrepancy with a 9.5 times lower affinity and a poor fit to the underlying model and was therefore rejected. The remaining 10 fragments were subjected to additional experimental characterization. An overview of these 10 fragments is given in **Table 1**. **Fig. 2** shows the sensorgrams and Langmuir regressions for four fragments of particular interest for the subsequent analyses (as described below).

**Figure 2.**
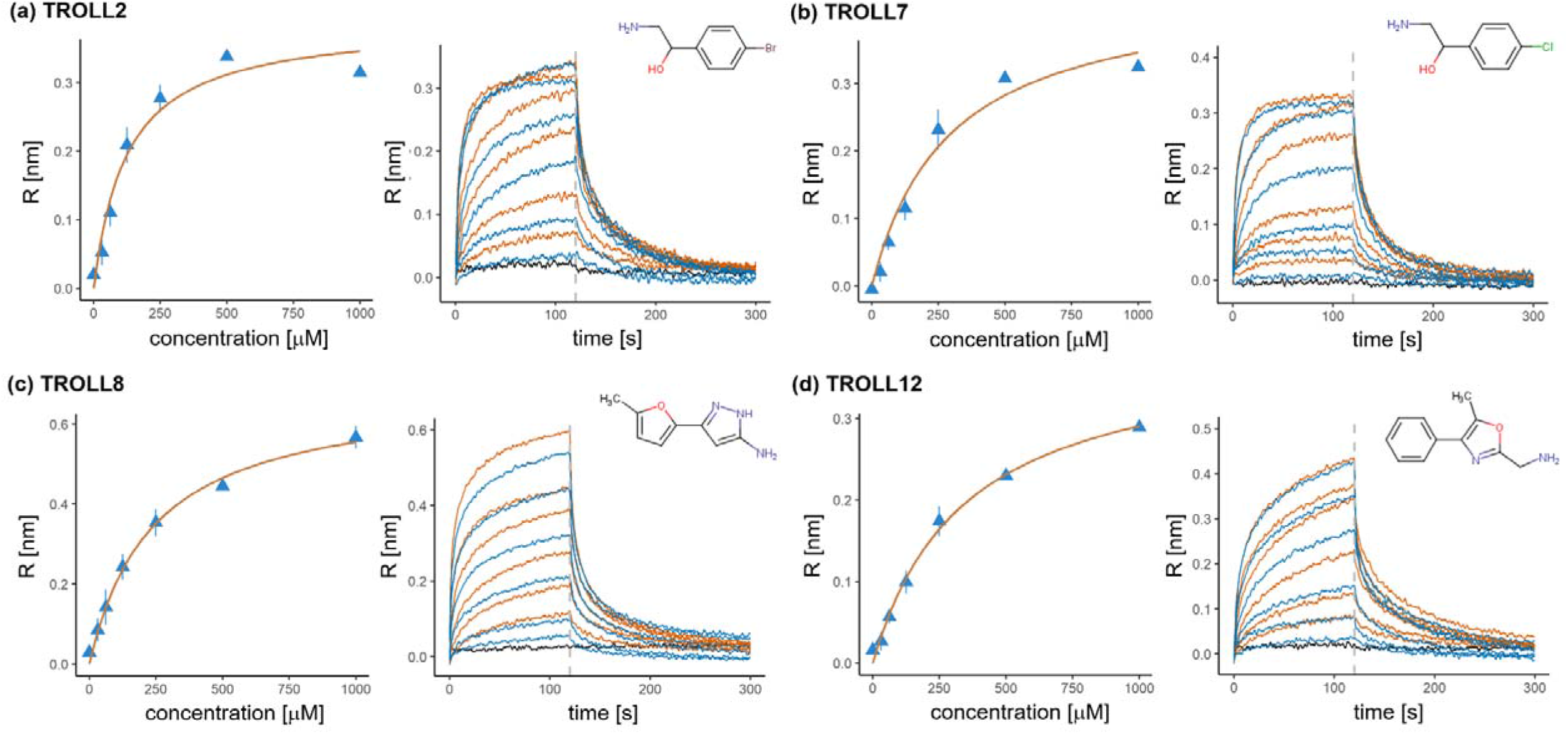
Sensorgrams and Langmuir models of selected fragments. Results of the dose-response analysis of selected N-domain hits. For each fragment the obtained shifts are plotted against the fragment concentration, with the fit to the Langmuir model shown in orange. Every concentration was measured twice and both values were used for fitting. Additionally, the sensorgrams of the fragments are shown after alignment and double referencing (first measurement: orange; second: blue). The negative control containing assay buffer only is shown in black. The dashed lines indicate the start of the dissociation phase. Sensorgrams which did not reach a plateau phase, because of unspecific binding or biphasic curve courses, received a penalty term in the calculated ScoreBLI (see method description in the Supporting Information).

**Table 1.**
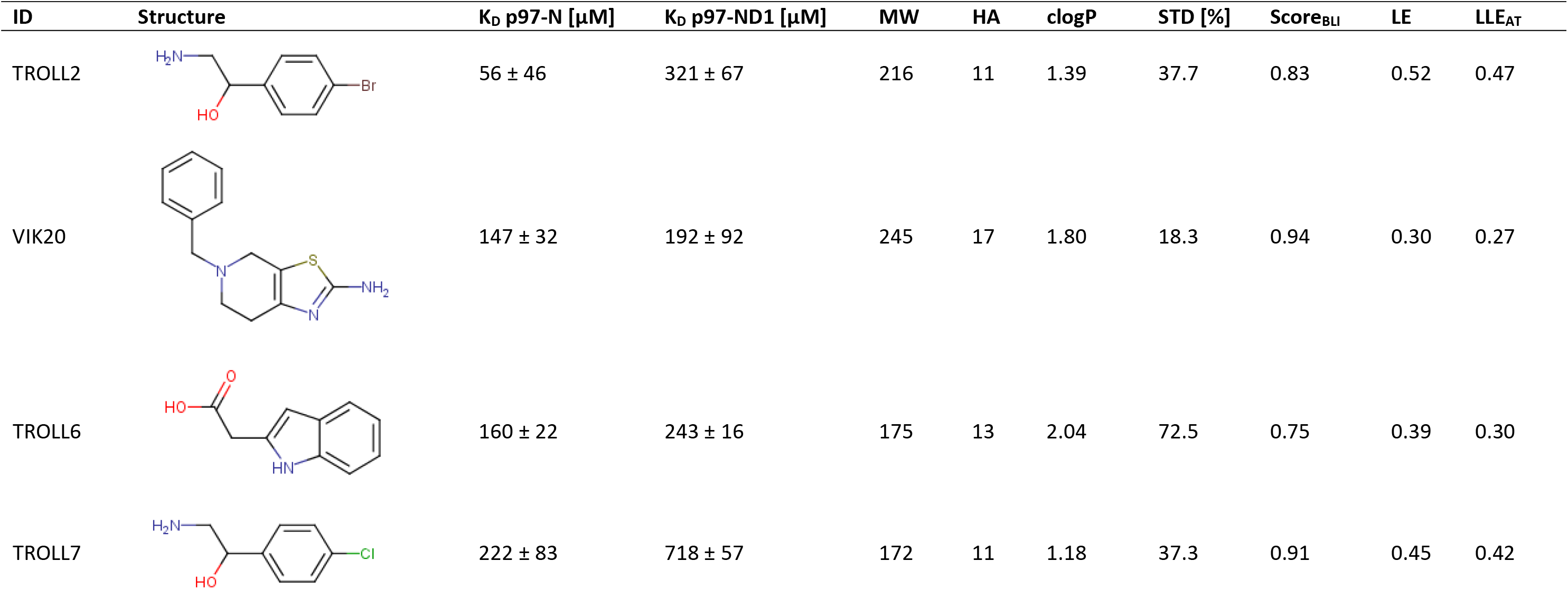
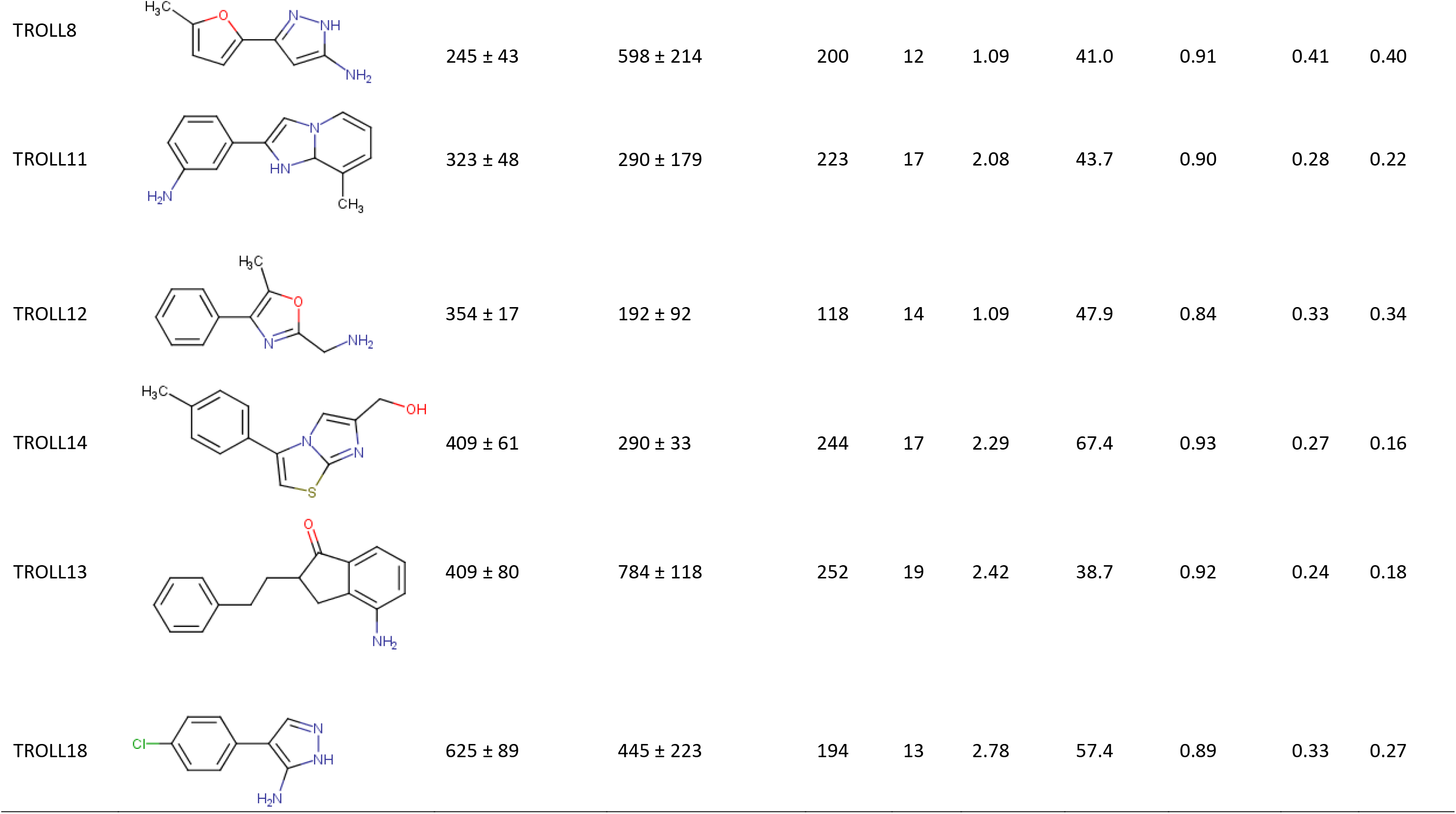
Overview of the identified fragments binding to the N-domain of p97. The compounds are sorted by their affinity for p97-N. The affinity obtained with the p97-ND1 construct is shown for comparison. Errors are derived from the fitting model. Molecular weight (MW), heavy atom count (HA) and calculated logP (clogP) are provided as descriptors. STD refers to the maximum STD signal measured with p97-ND1. The calculated ScoreBLI (based on the sensorgrams from measurements with p97-N) and two ligand efficiency metrics (LE and LLEAT, see discussion section for definition) are based on the affinities for p97-N. IDs are internal screening identification numbers.

| ID | Structure | K<sub>D</sub> p97-N [μM] | K<sub>D</sub> p97-ND1 [μM] | MW | HA | clogP | STD [%] | Score<sub>BLI</sub> | LE | LLE<sub>AT</sub> |
| --- | --- | --- | --- | --- | --- | --- | --- | --- | --- | --- |
| TROLL2 |  | 56 ± 46 | 321 ± 67 | 216 | 11 | 1.39 | 37.7 | 0.83 | 0.52 | 0.47 |
| VIK20 |  | 147 ± 32 | 192 ± 92 | 245 | 17 | 1.80 | 18.3 | 0.94 | 0.30 | 0.27 |
| TROLL6 |  | 160 ± 22 | 243 ± 16 | 175 | 13 | 2.04 | 72.5 | 0.75 | 0.39 | 0.30 |
| TROLL7 |  | 222 ± 83 | 718 ± 57 | 172 | 11 | 1.18 | 37.3 | 0.91 | 0.45 | 0.42 |

| ID | Structure | K<sub>D</sub> p97-N [μM] | K<sub>D</sub> p97-ND1 [μM] | MW | HA | clogP | STD [%] | Score<sub>B<sub>LI</sub></sub> | LE | LLE<sub>AT</sub> |
| --- | --- | --- | --- | --- | --- | --- | --- | --- | --- | --- |
| TROLL8 |  | 245 ± 43 | 598 ± 214 | 200 | 12 | 1.09 | 41.0 | 0.91 | 0.41 | 0.40 |
| TROLL11 |  | 323 ± 48 | 290 ± 179 | 223 | 17 | 2.08 | 43.7 | 0.90 | 0.28 | 0.22 |
| TROLL12 |  | 354 ± 17 | 192 ± 92 | 118 | 14 | 1.09 | 47.9 | 0.84 | 0.33 | 0.34 |
| TROLL14 |  | 409 ± 61 | 290 ± 33 | 244 | 17 | 2.29 | 67.4 | 0.93 | 0.27 | 0.16 |
| TROLL13 |  | 409 ± 80 | 784 ± 118 | 252 | 19 | 2.42 | 38.7 | 0.92 | 0.24 | 0.18 |
| TROLL18 |  | 625 ± 89 | 445 ± 223 | 194 | 13 | 2.78 | 57.4 | 0.89 | 0.33 | 0.27 |

### Chemical space analysis

To characterize the chemical properties of the fragments identified in this study, also in relation to those reported by Chimenti *et al.* [26], the positions of the fragments within the chemical space of the BiSS fragment library were determined. As a standard tool to analyze the chemical diversity of compound libraries [32,33,34], a principal component analysis (PCA) was applied. The PCA was based on 10 molecular descriptors (as specified in Materials and Methods). **Fig. 3** shows a plot of the first two principal components, which together describe 47.6% of the variance in the BiSS library. The positions of the previously reported fragments [26] and the newly identified compounds from the present study are mostly located in the same area in the lower right quartile of the plot: Eight of the 10 fragments from the present work and five of the six compounds previously reported can be found here. Only TROLL2 and its close analogue TROLL7, as well as ID2 are located in a different region of chemical space. The PCA analysis also indicates that there are additional enrichment options for further enhancements of screens against p97-N since the used fragment library does not fully coincide with the chemical space where most of the p97-N ligands can be found.

**Figure 3.**
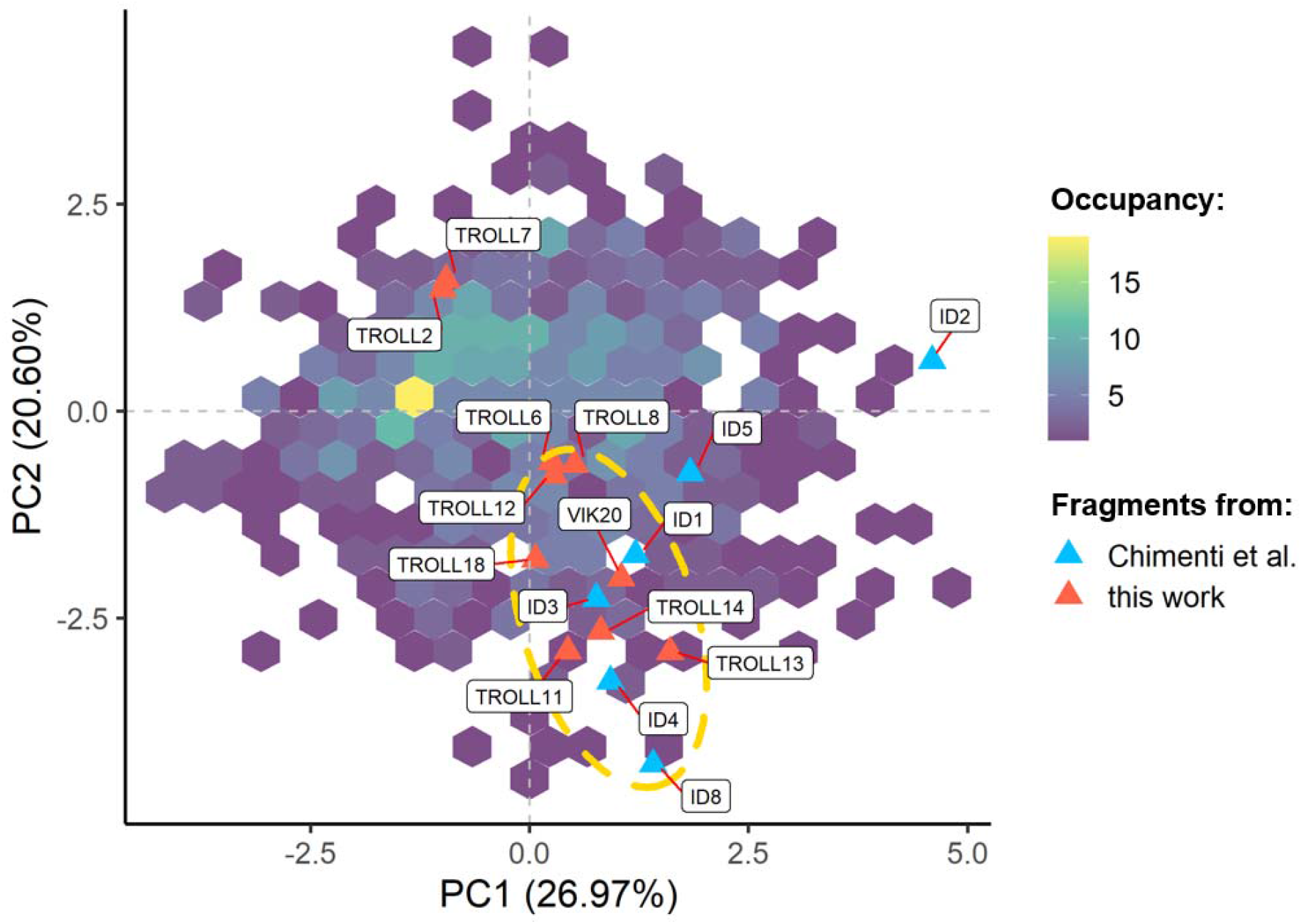
PCA analysis of the fragment library employed in this study. The first and second principal components together describe 47.6% of the variance in the dataset. The locations of the fragments identified in this screen (red triangles) and the fragments reported by Chimenti *et al.* [26] (blue triangles) are highlighted. The hexagons (colored purple to yellow) indicate how many compounds were available in the library for a particular region of the chemical space. The dashed orange ellipse highlights the region where most of the fragments of both screens are located, indicating that these fragments share similar physicochemical parameters.

### Ligand-based pharmacophore models

The fragments identified here as well as those reported earlier [26] were further analyzed for common or similar functional groups in terms of putative pharmacophore models. High structural agreement was found between TROLL12, TROLL14, VIK20 and ID4 regarding the position of an aromatic feature (i.e., a phenyl ring), an aromatic heterocyclic scaffold and the position of a hydrogen-bond donor function represented by a hydroxyl- or amino-group (**Fig. 4**, superpositions (a)-(c)). TROLL18 and ID2 share a 3-amino-pyrazole substructure in combination with a *para*-substituted phenyl ring (**Fig. 4, (d)**). The overlay of TROLL2 or its analogue TROLL7 with TROLL8 revealed two conserved hydrogen-bond donor functions, in combination with a hydrophobic, substituted aromatic ring system (**Fig. 4, (e)**). Furthermore, TROLL6 and ID3 display similar structural properties (**Fig. 4, (f)**). It can be hypothesized that the identified common features may be involved in interactions with residues in their p97 binding site.

**Figure 4.**
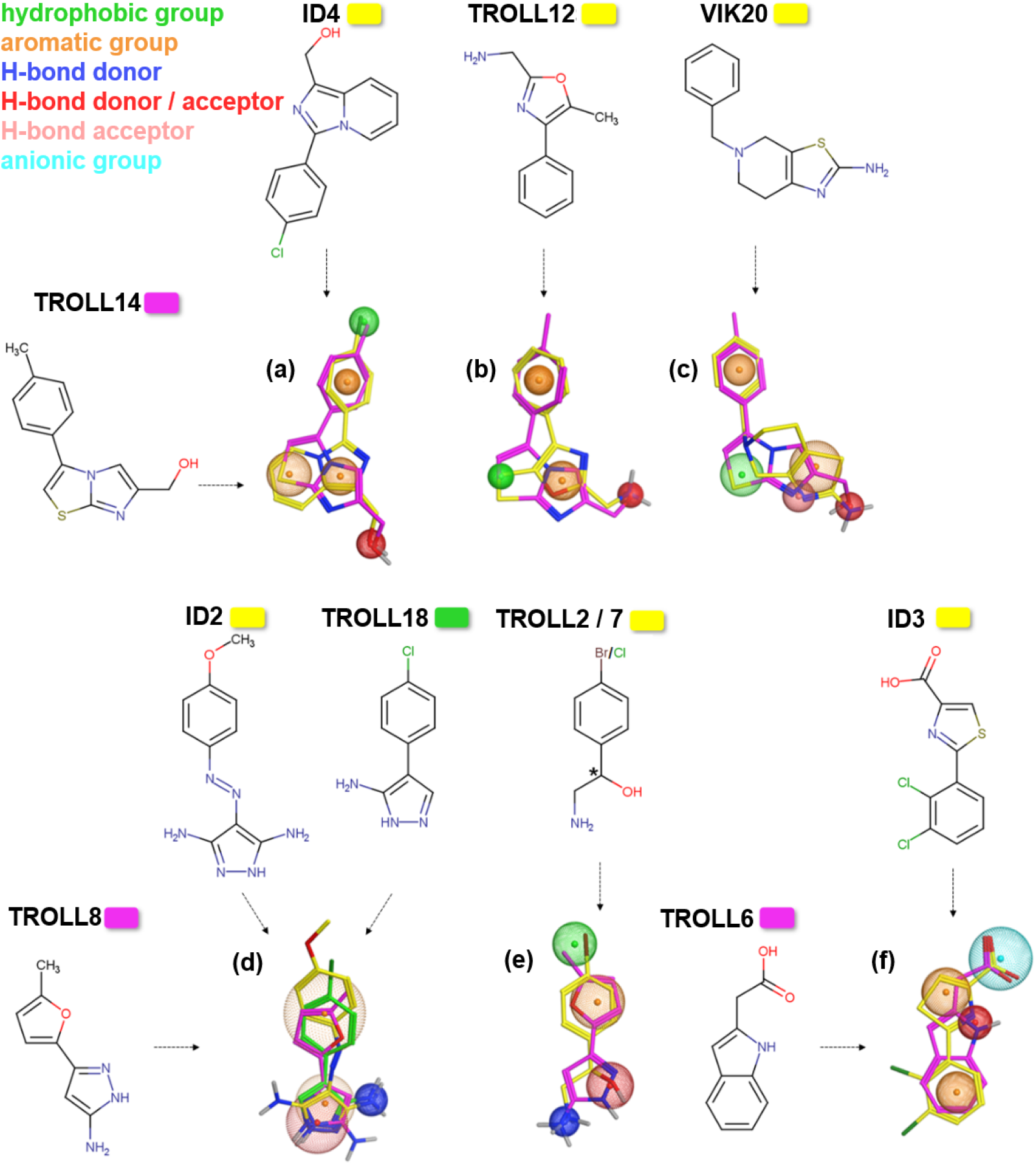
Pharmacophore models derived from selected fragments binding to p97-N. Fragments of this work and Chimenti *et al.* [26] were superimposed with respect to similar chemical properties or recognition elements. Spheres represent similar functional groups present in both fragments. The chemical properties of the spheres are explained in the colour code (top left).

### Ligand efficiencies of the identified fragments

One of the main questions in the early phase of drug discovery is the selection of compounds best suited for optimization towards lead structures. Besides an increase in affinity and specificity, especially for weakly binding molecules such as fragments, the optimization towards favorable physicochemical and ADME (Absorption, Distribution, Metabolism, and Excretion) profiles is of high importance. For this purpose, different ligand efficiency metrics were developed to guide the process of molecular optimization [35].

In this work, four different ligand efficiencies were calculated for the 10 selected molecules shown in **Table 1** based on their K_D_ value for p97-N, the heavy atom count and the clogP value: the ligand efficiency (LE) [36], the size-independent ligand efficiency (SILE) [37], the lipophilicity corrected efficiency (LELP) [38], and the modified lipophilic ligand efficiency by Mortenson and Murray (LLE_AT_) [39]. The four ligand efficiency metrics include indices for controlling the size (LE, SILE) and lipophilicity (LELP, LLE_AT_) of the fragments. For LE and SILE higher values indicate more promising candidates. A lower limit of 0.3 is commonly suggested for LE. LELP is calculated as (c)logP/LE and outlines the cost of the LE paid in lipophilicity; with a value of 0.3 for LE, LELP should lie between −10 and 10. The LLE_AT_ represents a more suitable metric for fragments, taking size and lipophilicity of the fragments into account. As with LE, its suggested lower cut-off is 0.3, and fragments with higher values are more promising [39]. The calculated indices were used to compare our fragment screen with screening campaigns reported in the literature. **Table 2** summarizes average values for selected physicochemical descriptors and ligand efficiency metrics of published hits from fragment screens [35] and compares them with the values for the fragments binding to the N-domain reported by Chimenti *et al.* [26] and the fragments identified in this study.

**Table 2.**
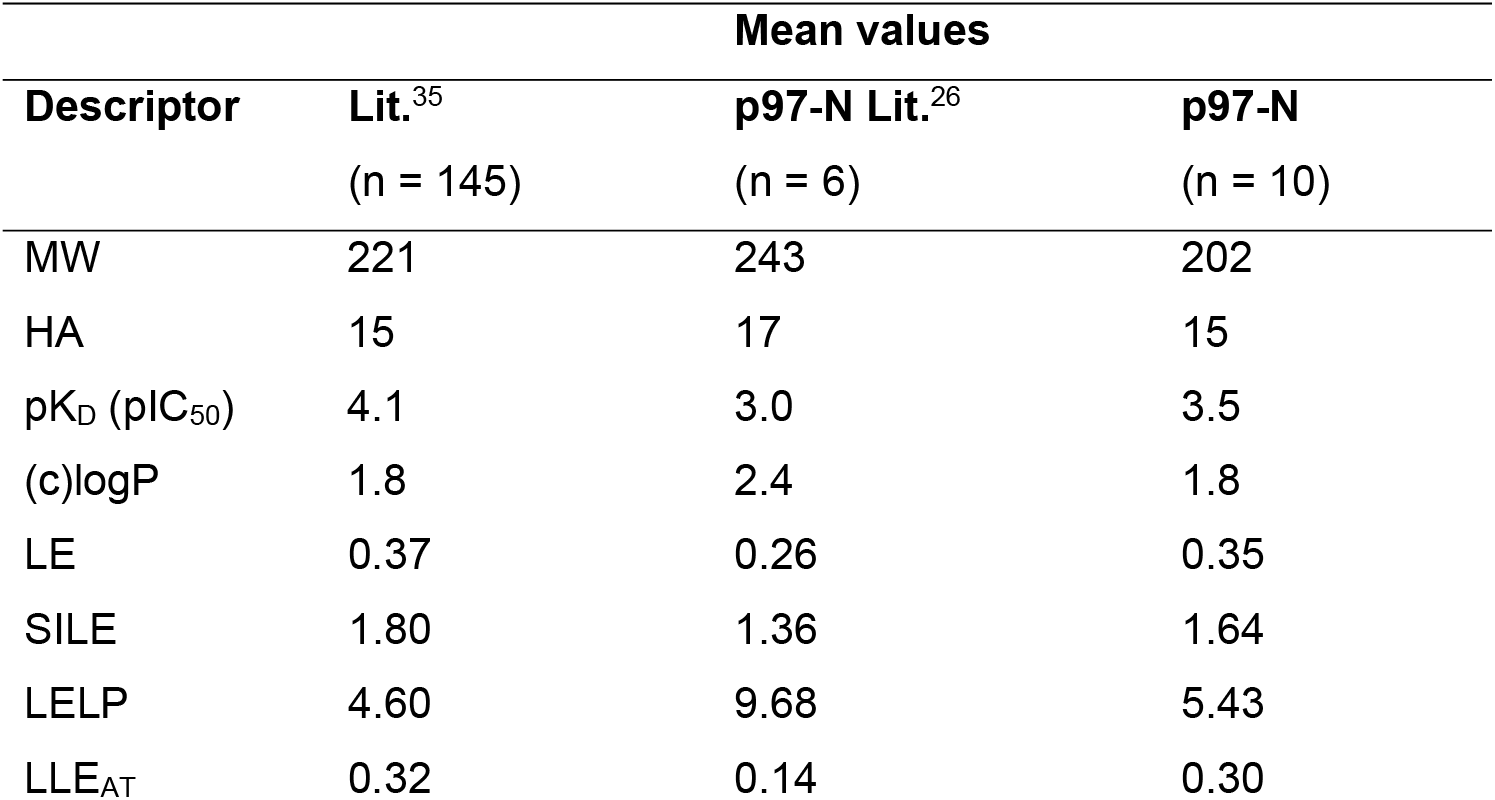
Comparison of descriptor values and ligand efficiency metrics for hits of fragment screens reported in the literature [35], hits of the p97-N screen reported by Chimenti *et al.* [26] and the hits of this work as listed in Table 1. MW is the molecular weight (in g/mol), HA the number of heavy (i.e., non-hydrogen) atoms; the four ligand efficiency indices are described in the text.

The average values for the 10 hits of our p97-N fragment screen are very close to the literature averages of successful fragment screens. In comparison to the fragments reported by Chimenti *et al.*, the newly identified compounds show superior properties. Especially the LELP and LLE_AT_ parameters (which are dependent both on clogP and affinity) indicate that binding of the fragments identified by Chimenti *et al.* may be dominated by hydrophobic interactions, which is not ideal for further optimization. The newly identified hits appear more promising because of a better balance between affinity and lipophilicity. In particular, the hit compounds TROLL2, TROLL7, TROLL8 and TROLL12 show an LLE_AT_ value above the cut-off of 0.3, which indicates that the binding is not dominated by hydrophobic interactions.

### Identification of p97 N-domain binding sites via mixed-solvent MD simulations

Mixed-solvent MD simulations are a common tool to analyze possible interaction hot spots for small molecules in protein binding sites [40–44]. In contrast to “blind” docking approaches where the protein is mostly rigid and no explicit solvent is used, mixed-solvent simulations provide a more realistic model of the protein in solution. Typically, fragments employed in this technology are even smaller in size than the fragments used in biophysical screens or the hits identified here. Nevertheless, Martinez-Rosell *et al.* [45] showed that the mixed-solvent simulation approach can handle such larger molecules as well. Consequently, mixed-solvent simulations were carried out to identify putative binding sites. The approach used here is based on the “Site Identification by Ligand Competitive Saturation” (SILCS) methodology [40], where a high fragment concentration is subjected to an all-atom explicit-solvent MD simulation. The aggregation of fragments is prevented by dummy atoms that are placed into the molecules to exert a repulsive force between the fragments.

We simulated each fragment at a concentration of 1 M and calculated occupancies of the fragments on the surface of p97-N to identify possible binding regions. For every fragment, the two sites with the highest occupancies were selected, and the binding site was defined by all amino acids within 6 Å of the fragment’s dummy atom. Most of the residues identified by this approach are in the Nc-subdomain of p97-N, especially in the area of the SHP binding region (**Fig. 6**), while only a few residues are located in the Nn-subdomain. This suggests the Nc-subdomain and particularly its SHP-binding region as the most “druggable” part of the N-domain. Based on the observations in the mixed-solvent simulations, the SHP binding region was subdivided into three regions (SHP I, SHP II and SHP III). SHP I is the region addressed by the conserved leucine residue of the SHP binding motif. SHP II is addressed by the aromatic amino acids F228 of UFD1 and W242 of Derlin-1 [46,47]. SHP III is an adjacent hydrophobic cavity not directly addressed by SHP motif containing cofactors, which harbors a solvent-exposed cysteine residue (C184).

**Figure 6.**
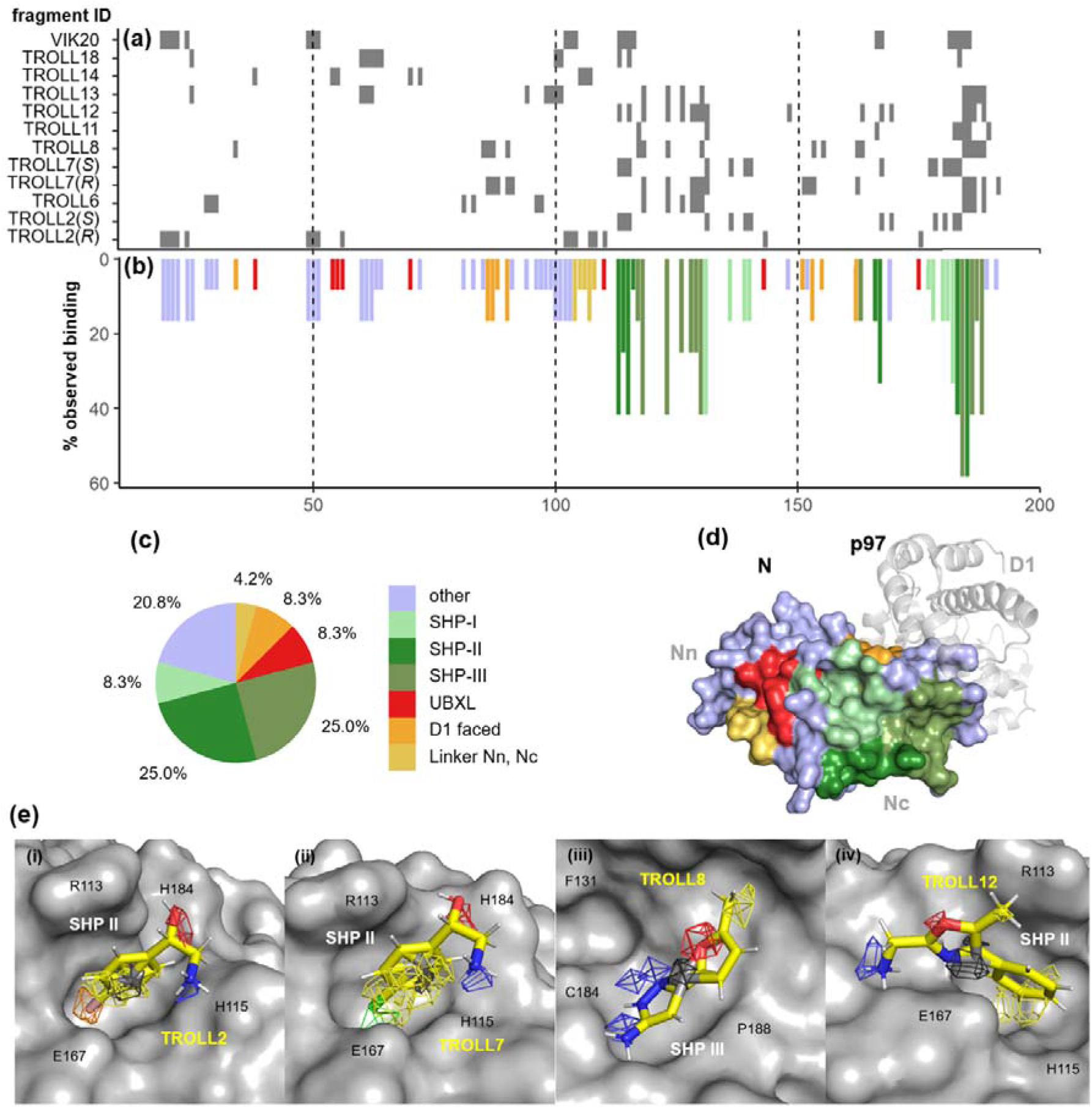
Results of mixed-solvent MD simulations. The 10 hit fragments of Table 1 were simulated, including both enantiomers of TROLL2 and TROLL7 in separate simulations. **(a)** Binding-site residue map. Grey rectangles specify amino acids along the primary sequence (horizontal axis) which are in the vicinity (< 6 Å from the dummy atom) of the two sites with highest occupancies of the corresponding fragment. **(b)** Relative propensity of observed binding interactions. The bars indicate in how many of the 12 individual simulations the residue was identified as binding-site residue according to panel (a), using the color code of panel (c). **(c)** Pie chart with the distribution of the two sites with highest occupancy for each fragment across the N-domain surface (N=24). **(d)** Location of the corresponding regions on the surface, color-coded according to the pie chart. **(e)** Illustration of fragments for which a binding mode could be postulated from the mixed-solvent simulations: **(i)** TROLL2 and **(ii)** TROLL7 in the SHP II site; **(iii)** TROLL8 in a cavity near the SHP binding region (SHP III); and **(iv)** TROLL12 in SHP II, in a different orientation compared to TROLL2 and TROLL7.

Within the identified binding sites, the occupancies of the heavy atoms of each fragment were further inspected to obtain information about the orientation of the molecules and to distinguish between specific and unspecific binding (**Fig. 6(e)**). In case of the four fragments TROLL2, TROLL7, TROLL8 and TROLL12, the resulting occupancies indicated an orientational preference, thus allowing to predict a binding mode based on the observed densities. For TROLL2 (which was simulated as both the *R*- and the *S*-enantiomer) only the *S*-enantiomer was predicted to bind within a pocket formed by residues R113, H115, E167 and H183 (SHP II, **Fig. 6(e, i)**). The closely related analogue TROLL7 presumably exhibits the same interactions (**Fig. 6(e, ii)**). TROLL12 was also found in this region, but differently oriented (**Fig. 6(e, iv)**), whereas TROLL8 bound to a sub-pocket near C184 in the SHP III region (**Fig. 6(e, iii)**). Additional occupancies for the methylfurane ring of TROLL8 were also identified in the SHP II region.

### Crystal structure of TROLL2 in complex with the N-domain of p97

To confirm the predicted binding sites, X-ray crystallographic studies were performed. As the fragment with the highest binding affinity in the BLI screen, a high-resolution structure of TROLL2 in complex with p97-ND1 could be obtained. The identification of the Br-containing TROLL2 fragment and its binding site was assisted by the anomalous signal of the halogen atom. The structure was solved by molecular replacement using PDB structure 5DYG as the search model. In analogy to Tang & Xia, the same hexagonal space group (P622) with one ND1-monomer in the asymmetric unit was obtained [48].

In four data sets a significant anomalous signal corresponding to one binding site as determined by the SHELXCD pipeline [49] could be identified. To confirm that the anomalous signal is not an artefact of the crystallization conditions, data from apo-crystals (n=43) obtained under identical conditions without any fragment were analyzed for anomalous signals. No anomalous signal could be identified in the same region, and any observed anomalous signal was significantly weaker by 6σ compared to the TROLL2 data. These data demonstrate that the observed anomalous signal is due to the presence of the TROLL2 fragment. The data set with the strongest anomalous signal was refined to a crystallographic R-factor of 19.20% and a free R-factor of 23.53% at 1.73 Å resolution with Phenix [50] (cf. **Table S1** for data collection and refinement statistics). A superposition of PDB structure 5DYG with our structure in complex with TROLL2 showed a root mean square deviation (RMSD) of only 0.2 Å for the C_α_-atoms.

The anomalous signal of the Br-atom was unambiguously assigned to the binding pocket identified in the mixed-solvent simulations (**Fig. 7 (a)**). A polder omit map contoured at 3σ showed a clear density for the Br-atom and parts of the neighboring carbon atoms of the phenyl ring (**Fig. 7 (b)**). The hydrophobic substructure of TROLL2 interacts with a region formed by the aliphatic parts of residues R113, H115, E167 and E185. The phenyl ring of TROLL2 adopts a favorable orientation for a T-shaped interaction with H183. Due to weak electron density the interactions of the hydroxyl and amino groups of the inhibitor remain undefined. The binding region contains a contact from an arginine (R113*) of a symmetry-related N-domain which is not part of the p97-ND1 hexamer. This arginine engages in a cation-π interaction with the phenyl ring of TROLL2. A detailed comparison of the results of the mixed-solvent MD simulations shows a high level of agreement in the orientation of the resulting occupancy densities with the experimentally obtained structure. Additionally, the simulations revealed that R113 from the same monomer can also interact with the phenyl ring of the fragment via a cation-π interaction; in solution, it should, thus, be able to replace the R113* interaction resulting from crystal packing.

**Figure 7.**
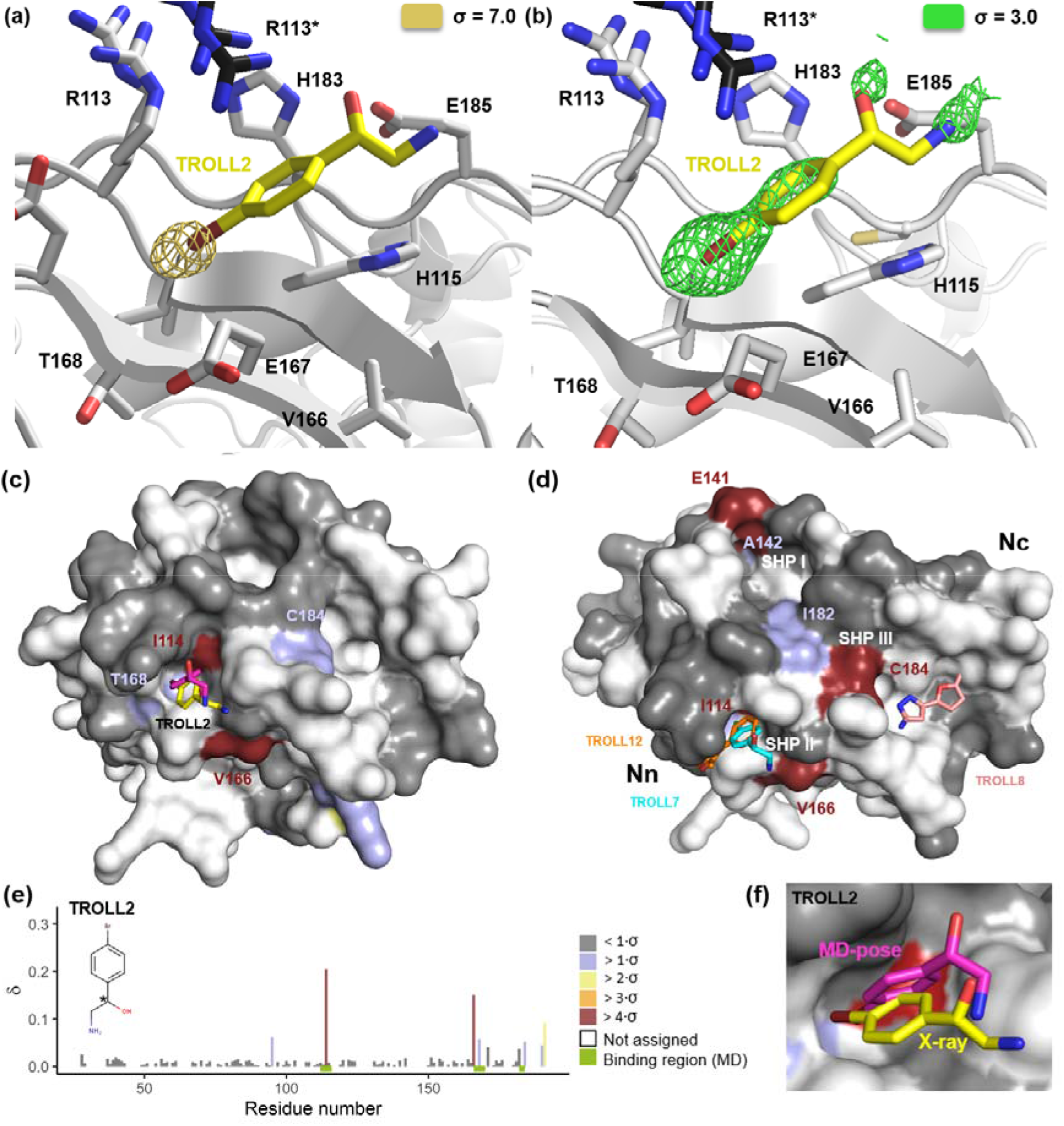
Crystal structure of TROLL2 in complex with the SHP II binding site of p97-N and binding sites identified via ^1^H-^15^N-HSQC NMR measurements. **(a)** Anomalous map contoured at a sigma level of 7 highlighting the single signal in the asymmetric unit, which harbors one ND1 protomer. Only the binding region superimposed with the final refined model is shown, with TROLL2 carbon atoms highlighted in yellow (bromine atom in dark red). **(b)** Polder omit map of TROLL2 calculated with Phenix at a sigma level of 3. **(c)** Chemical shift changes mapped onto the surface representation of p97-N for TROLL2, following the color code defined in panel (e). **(d)** Maximal chemical shift changes from either TROLL7, TROLL8 or TROLL12 mapped onto the surface representation of p97-N (cf. **Fig. S6** for individual representations for each fragment). The additionally shown binding poses in stick representation in (c) and (d) were obtained from the mixed-solvent MD simulations, except for the TROLL2 pose shown in yellow, which corresponds to the crystal structure. **(e)** Chemical shift changes observed for TROLL2 plotted against the primary sequence. **(f)** Close-up view of the SHP II region with MD-pose in magenta and the crystal structure in yellow. The surface of I114, for which the strongest chemical shift was observed, is shown in red.

### 1H-15N-HSQC NMR measurements with the N-domain of p97

To clarify whether the binding site of TROLL2 obtained in the crystal structure is due to the symmetry-related N-domain crystal contact and since no crystal structures of p97-ND1 in complex with TROLL7, TROLL8 and TROLL12 could be obtained, ^1^H-^15^N-HSQC NMR measurements with the isolated N-domain of p97 were performed to further complement the mixed-solvent MD simulations.

The ^1^H-^15^N-HSQC NMR experiments of all fragments showed chemical shift changes within the regions postulated by the mixed-solvent MD simulations (**Fig. S6**). Specifically, the largest chemical shift changes could be observed for TROLL7 and TROLL2 for residues I114 and V166, which are both located in the SHP II region. Small shifts for both fragments were also detected for T168 within the SHP II and C184 in the neighboring SHP III region. Additional chemical shifts were assigned to R95, K190 and R191. In full-length p97 these regions are in contact with the D1-domain and not accessible or exhibit a different conformation. Due to this fact the observed chemical shifts of these amino acids were not further considered. The remaining chemical shift changes were all in or in close proximity to the SHP II region and in agreement with the binding region identified by the mixed-solvent MD simulations and in the crystal structure with TROLL2. TROLL12 only showed for I114 a slight chemical shift, originating within the predicted region of the mixed-solvent MD simulations, but the data here were not as robust as for TROLL2 or TROLL7.

For TROLL8 a significant chemical shift was observed for C184 in the SHP III region (predicted by the simulations) and for E141. E141 and A142 are located between the SHP I site and the UBX/L binding region. Both regions were not identified as possible binding sites by the mixed-solvent MD simulations; however, because of the lack of a suitable pocket in the UBX/L region, a binding to the SHP I region seems to be more likely. Additional minor chemical sifts were located again within the SHP II region. The simulations showed here a low occupancy for the compound’s methylfurane ring.

The observed chemical shift changes were not particularly strong; nevertheless, they only occurred in the presence of the fragments and not in the negative controls under identical conditions. The chemical shifts were also remarkably focused to single amino acids. It should also be noted that parts of the SHP II and most of the SHP III sites could not be assigned and so no experimental data were available for these areas. Titration experiments for affinity determination were technically not possible due to the poor solubility of the fragments. Interestingly, no strong chemical shift changes were observed within the N-terminal subdomain, in agreement with the general outcome of the mixed-solvent MD simulations. In summary, the results of the ^1^H-^15^N-HSQC NMR measurements were in good overall agreement with the MD simulations and the crystal structure of the TROLL2-p97 complex and confirmed the addressability of the SHP binding site with small molecules (cf. **Fig. 7 (c)-(f)**).

## Discussion

### BLI is a well-suited method for fragment screening

So far the BLI method has been reported only sporadically in the field of fragment-based drug design [27]. Here, we established a BLI assay as a robust and well-performing fragment-screening platform. Key advantages of BLI as a fluid-free system are an easier setup and handling in comparison to other biosensor platforms. The present study demonstrates that this method is a suitable biophysical approach for detecting the binding of low molecular-weight compounds, thus providing promising candidates for a fragment-based drug design approach, also in the context of PPI inhibitors.

The N-domain of p97 as main cofactor interaction site and potential target for PPI inhibitors was the focus of this work. Due to the absence of a specific binder as positive control for the isolated N-domain, the screening quality was first assessed with the p97-ND1 construct. The screen on p97-ND1 revealed a Z-factor of 0.67 and a signal distribution that reflects a high overall quality of the approach. The optimized screening conditions for the ND1 domain were then transferred to the isolated N-domain. An overlap of 18 fragments between the two screens was detected, where most of the N-domain hits were also identified with the ND1 construct (see **Table S2**). Stringent screening allowed to identify 10 high-confidence hits, which were subsequently confirmed by STD-NMR and additional BLI measurements using the ND1 construct. The satisfying performance of the screen was further demonstrated when comparing our results with the SPR-based screen of the ND1 domain conducted by Chimenti *et al.* [26] using a principal component and pharmacophore analysis. The analysis showed that the BLI-based screen was capable of identifying similar fragments in terms of their position in the chemical space, but also revealed new binders exhitibing different properties.

Nonetheless, even when promising fragment candidates have been identified for a target protein, the selection of candidates for a more detailed experimental characterization and further optimization is a challenging task [51]. The Score_BLI_ applied in this work proved helpful for eliminating hits with unwanted properties, such as unspecific binding. With respect to ligand efficiencies the fragments identified here display superior properties compared to those reported earlier [26]; in particular, our hits are not dominated by hydrophobic interactions and exhibit more favorable ligand efficiencies for subsequent optimization. The use of mixed-solvent MD simulations for the further selection of fragments binding to specific regions of the target protein proved to be a highly beneficial step as the postulated binding sites could be confirmed by X-ray and ^1^H-^15^N-HSQC NMR measurements. Taken together, this BLI-based fragment screening strategy in combination with subsequent *in silico* tools turned out to be a highly successful approach.

### Identification of starting points for novel PPI inhibitors targeting the p97 SHP-motif binding site

A selective inhibiton of cofactor binding to the N-domain of p97 would represent a new and valuable avenue in the field of p97 inhibitors. The work described here represents a significant advance towards this goal. The structural relevance of the identified fragments TROLL2, TROLL7, TROLL8 and TROLL12 as starting points for the development of a PPI inhibitor is shown in **Fig. 8**. A comparison of the binding sites in the crystal structures of the SHP-motif containing cofactors Derlin-1 and UFD1 with the identified target sites of the fragments reveals a high degree of overlap, in particular in the SHP-II region. The identified fragments address two important hotspots of the PPI of SHP-motif containing cofactors: (i) The postulated binding poses of TROLL7 and TROLL12 derived from the mixed-solvent MD simulations as well as the binding pose found in the crystal structure of TROLL2 mimic the binding of the aromatic amino acids F228 of UFD1 and W242 of Derlin-1 in the SHP-II region, respectively. The importance of this hotspot interaction for the binding of the SHP-motif containing cofactor UFD1 is demonstrated in the reduced affinity for the F228A mutant of p97 [46,52]. (ii) Furthermore, the ^1^H-^15^N-HSQC NMR measurements indicated that TROLL8 binds adjacent to the SHP-III region into a sub-pocket referred to as SHP-I region, which is addressed by L235 (UFD1) or L248 (Derlin-1). Like the aforementioned aromatic residues these aliphatic side chains are hotspot residues and are important for the binding to p97 as shown by their respective replacement with alanine [52,47].

**Figure 8.**
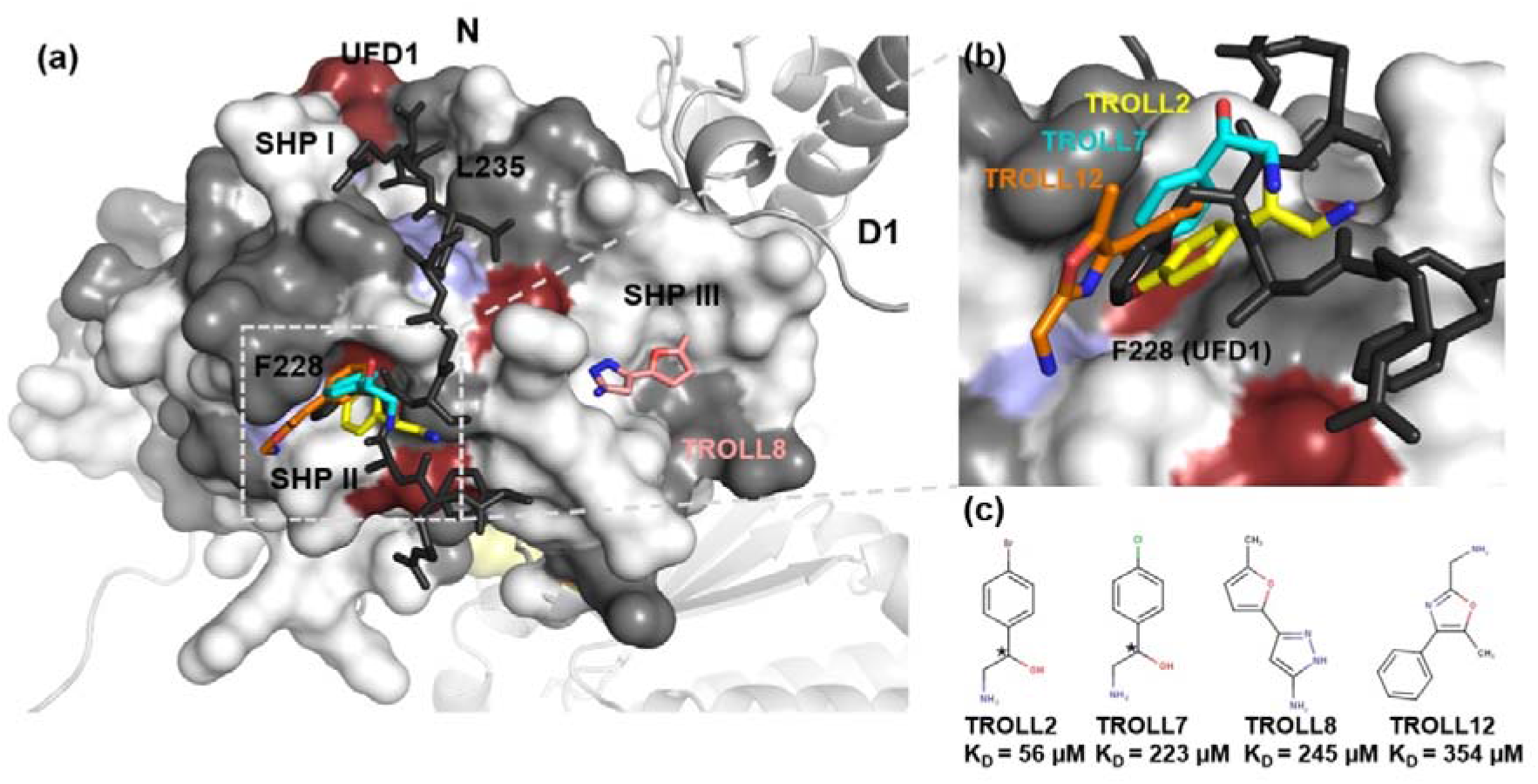
Overlap between postulated and experimentally verified fragment binding sites on p97 with the interaction sites of UFD1. **(a)** Binding sites of TROLL7, TROLL8 and TROLL12 identified in the mixed-solvent MD simulations, as well as the crystal structure of TROLL2 superimposed with the bound UFD1-SHP-peptide (PDB entry 5B6C) mapped onto a surface representation of the N-domain. Additionally, the highest chemical shifts of the ^1^H-^15^N-HSQC-NMR measurements are shown colored in terms of standard derivation as in Fig. 7. **(b)** Zoom into the SHP II binding site, which is occupied by F228 in the structure of the p97N-UFD1 peptide complex and is addressed by TROLL2, TROLL7 and TROLL12. Based on the ^1^H-^15^N-HSQC-NMR measurements, and in contrast to the findings from the mixed-solvent MD simulations, TROLL8 seems to bind to a sub-pocket (SHP I) which is occupied by L235 in the structure of the p97N-UFD1 peptide complex. **(c)** Structures and affinities of the characterized fragments binding to p97-N.

Taken together, our results suggest that the design of a PPI inhibitor for the aforementioned binding site should be possible, since the SHP-motif binding region on p97 has the ability to bind to small molecules. Besides the potential to more comprehensively understand the regulation of p97 by its cofactors, a PPI inhibitor targeting the interaction between p97 and SHP-motif containing cofactors would yield a promising strategy to modulate the degradation of ubiquitinated proteins, with the ultimate promise of a new strategy to treat cancer.

This successful fragment screen against the N-domain of p97 used the previously underemployed BLI approach and establishes it as a viable alternative to SPR. Importantly, the characterization of the obtained hits resulted in the first crystal structure of a small molecule in complex with the SHP-motif binding region of p97. Additional binding sites predicted by mixed-solvent simulations could be confirmed by ^1^H-^15^N-HSQC-NMR measurements, and a high agreement of different biophysical methods on the binding of the top-ranked fragments was achieved. The compounds identified in this study display superior physicochemical properties for further drug design and provide promising starting points for the development of a PPI inhibitor addressing the SHP-motif binding site in the N-domain of p97.

## Materials and methods

### Cloning, Protein Expression and Purification

Constructs of p97-N (aa 21-199, N-terminal 3C protease-cleavable His_6_-tag) and p97-ND1 (aa 2-481, N-terminal TEV protease-cleavable His_6_-tag) harbouring a C-terminal Avi-tag were generated by inverse mutagenesis as described [28]. The correct nucleotide sequence of all constructs was verified by DNA sequencing (Microsynth Seqlab, Göttingen, Germany). Expression and purification were carried out as described in Hänzelmann *et al.* [53]. Biotinylation of Avi-tagged p97-N and p97-ND1 was conducted as described [28] followed by size exclusion chromatography.

For crystallization p97-ND1(L198W) (residues 1–460) encoding a C-terminal His_6_-tag with a two amino acid linker (RS) according to the construct used by the Xia lab [48] was generated by the restriction free cloning method [54] and site-directed mutagenesis (QuickChange Site-Directed Mutagenesis Kit, Agilent). Expression and purification were carried out as described in Tang and Xia [48].

Bacterial expression of ^15^N-labelled protein was performed according to a published protocol [55]. p97N (aa 1-208, N-terminal TEV protease-cleavable His_6_-tag [56]) was expressed in *E. coli* BL21 (DE3) RIL cells. Cells were grown in LB medium until the OD_600_ reached a value between 0.6 and 0.7. After centrifugation, the medium was removed and the cells were resuspended in M9 minimal medium supplemented with ^15^NH_4_Cl as the sole nitrogen source, where the volume of the M9 medium corresponded to one quarter of the amount of LB medium previously used. After incubating the cells for 1 h at room temperature, p97 expression was induced with 1 mM IPTG overnight at 16°C. The protein was purified in a buffer containing 50 mM Tris pH 8, 150 mM KCl, 5% glycerol (v/v), 5 mM MgCl_2_ and 5 mM β-mercaptoethanol by Ni-IDA affinity chromatography and size exclusion chromatography (Superdex 200, Cytiva). Prior to size exclusion chromatography, the protein was dialyzed overnight at 4°C in the presence of TEV protease (1:50) followed by Ni-IDA affinity chromatography to remove uncleaved His-tagged protein.

### BLI fragment screening

#### Fragment library

For fragment screening, the library compiled by the Facility for Biophysics, Structural Biology and Screening at the University of Bergen (BiSS) was used. This fragment library is a 679 compound subset of the OTAVA chemicals solubility library which was filtered to exclude similar compounds and compounds with unwanted functionalities [57]. The fragments have a mean molecular weight of 202 Da and an average clogP (calculated partition coefficient) of 1.52. The fragments contain on average two ring systems, one hydrogen-bond donor, two hydrogen-bond acceptors and two rotatable bonds. Hits from the initial screenings were ordered from OTAVA chemicals for further experimental characterizations.

#### BLI sensor preparation

Protein constructs for BLI measurements were enzymatically biotinylated using BirA [28]. Biotinylated p97 was loaded on Super Streptavidin Biosensors (SSA) obtained from Sartorius as follows. SSA-sensors were equilibrated in PBS-buffer, loaded with a protein solution containing 50 µg/mL (p97-ND1) or 100 µg/mL (p97-N) protein, blocked with biocytin and washed in PBS buffer. Reference SSA-sensors were set up by blocking them with a 10 µg/mL biocytin solution for 5 min [27]. The SSA-sensors were loaded to a shift height of up to 7 nm with p97-ND1 and 10 nm with p97-N, respectively.

#### General screening setup

Screening plates were prepared by adding 10 µL of a 10 mM DMSO stock solution of library compound to 190 µL of PBS buffer (containing 0.05% TWEEN 20, 1 mM DTT and 5 mM MgCl_2_ [only for p97-ND1]) in 96 well plates (Greiner). The final DMSO concentration was 5% (v/v). All BLI measurements were performed on an Octet RED96e (Sartorius) device.

All fragments were measured twice at a concentration of 500 µM by screening every plate in the forward and backward direction. The assay settings were as follows: Baseline measurement 15 s; association time 60 s and dissociation time 300 s. The resulting data were processed using the double reference method of the Octet Analysis software.

#### Initial screening using the ND1-domain

Signals of positive (500 nM ADP) and negative controls (buffer) were collected before and after each screening plate. The signals of each sensor were corrected for partial inactivation of the protein over time and for different protein loading states of the sensors using the signals of the 500 nM ADP positive control (see **Fig. S7** for details on the correction approach).

#### Initial screening using the N-domain

One fragment (ID5) reported to bind to the N-domain [26] could be purchased from Enamine and was tested as a positive control. The compound showed an uninterpretable behavior in the BLI assay using the ND1- as well as the N-domain and no valid dose response could be obtained. Because of the lack of another valid positive control (small molecule) for the isolated N-domain of p97, the assay conditions of the ND1 screen were transferred to the N-domain screen. Every sensor with immobilized p97-N was used for measuring only two screening plates and then discarded. Sensors showing artificial signals during the screening were replaced. The signals were normalized against the protein loading signal of each sensor. Signals with artificially high or large negative values based on visual inspection were eliminated.

#### Concentration dependencies

Selected fragments of the initial screenings were confirmed by dose-response-titrations using six concentrations ranging from 1000 to 31.3 µM in a 1:1 dilution series. Every concentration was measured twice. The assay settings were as follows: baseline measurement 15 s; association time 120 s and dissociation time 180 s. The double referenced data were further analyzed in Origin Pro to estimate the affinity by fitting the signals of the steady states to a 1:1 Langmuir model. For the fitting all measured data points (n=2) were considered.

### Principal component analysis (PCA) and ligand-based pharmacophore models

The PCA for fragments of the BiSS library was carried out in R using the *prcomp()* function. It was based on the following descriptors: HA, number of heavy atoms; HDon, number of hydrogen-bond donors; HAcc, number of hydrogen-bond acceptors; TPSA, topological polar surface area; Nrot, number of rotatable bonds; NChir, number of chiral atoms; NRings, number of ring systems; clogP, calculated logarithmic partition coefficient; Narom, number of aromatic atoms; Fsp3, fraction of sp3 hybridized atoms. All descriptors were calculated with MOE (Molecular Operating Environment (MOE), 2019.01, Chemical Computing Group ULC, 1010 Sherbooke St. West, Suite #910, Montreal, QC, Canada, H3A2R7, 2021). A flexible molecular alignment was carried out in MOE with the selected fragments of this study and the fragments reported by Chimenti *et al.* [26]. The aligned molecules were analyzed using the 3D pharmacophore builder in MOE.

### Mixed-solvent MD simulations

For the mixed-solvent MD simulations the complex structure of p97-N with the SHP-motif of UFD1 (PDB:5B6C) [46] was used. All water and buffer molecules as well as the UFD1 peptide were removed. The structure was prepared and protonated (at pH 7) in MOE using default values. For mixed-solvent simulations the Site Identification by Ligand Competitive Saturation (SILCS) method [40] was used. To prevent aggregation of lipophilic fragments, dummy atoms were inserted at the center of mass of each ligand. These atoms served as virtual sites for repulsive interactions (only between fragments) using the Lennard-Jones parameters adopted from the SILCS method (*ε* = −0.01 kcal/mol; *R_min_* = 24.0 Å), but without applying a switching function. 1.14*CM1A-based charges for the ligands were retrieved from the LigParGen web server [58–60]. A cubic simulation box with a 1 M fragment concentration and 5 nm edge length was solvated with TIP3P water. The solvation box was energy-minimized for 50,000 steps using the steepest descent minimization, equilibrated under NVT and, subsequently, NPT conditions for 100 ps each, and then simulated for 1 ns (NPT) using GROMACS [2019.01] [61] with the OPLS-AA force field [62] in order to obtain a mixed-solvent box with a homogenous ligand concentration. Coupling of temperature and pressure was performed using the velocity rescaling thermostat and the Parrinello-Rahman barostat [63]. The protonated protein structure was then solvated with this mixed-solvent box, resulting in a cubic starting system with at least a 1 nm distance from the simulation box border to the protein. The system was again minimized and equilibrated as described above and then subjected to a 40 ns production run with 2 fs time steps, writing out snapshots every 10 ps. Positional restraints on protein heavy atoms were applied during the equilibration phase. Four replicas were performed for each ligand, resulting in a total simulation time of 160 ns per fragment. For TROLL2 and TROLL7 simulations of both stereoisomers were carried out. For TROLL2, TROLL7, TROLL8 and TROLL12 the simulations were extended to a total simulation time of 400 ns. Afterwards, the trajectories were aligned on the C_α_ atoms and occupancy densities were calculated for the dummy atoms and selected heavy atoms of the fragments with a resolution of 1.0 Å using the volmap plugin in VMD [64]. The densities of individual replicas were added using the function volutil resulting in a sum density map for each fragment.

### Crystallization

The p97-ND1 L198W variant was incubated with ADP (molar ratio 1:13) and crystals were grown at 20 °C by vapor diffusion in hanging drops containing equal volumes (1:1) of protein solution (15 mg/ml) and reservoir solution consisting of 5% PEG 600 (v/v), 4.0 M sodium formate (pH 6.0), 5% glycerol (v/v) and TROLL2 in a final molar ratio of 1:10.

### Data Collection and Structure Determination

p97-ND1 crystals were cryo-protected by soaking in mother liquor containing increasing amounts (10%-30% [v/v]) of glycerol and TROLL2 in a molar ratio of 1:10. The crystals were flash-cooled in liquid nitrogen, and data collection was performed at 100 K at the ESRF in Grenoble (beamline ID30B, wavelength 0.9184 Å). Data were processed using XDS [65] and STARANISO [74]. The anomalous signal of the bromine atom was detected using SHELX [49] and AnoDe [66]. Further calculations were performed using the CCP4 Suite [67] and Phenix [50]. Phases were obtained by molecular replacement using Phaser with the ND1-domain of p97 (PDB: 5DYG [48]) as search model. TROLL2 was modelled based on the difference map. The structure was ultimately refined in Phenix [50]. TLS groups were obtained from the TLSMD-Webserver [68]. Restraints for TROLL2 were calculated using eLBOW in Phenix.

### STD-NMR spectroscopy

All 1D ^1^H STD NMR spectra [31] were collected on a Bruker Avance spectrometer with a magnetic field strength of 9.4 T, corresponding to a ^1^H resonance frequency of 400.13 MHz. The spectrometer was equipped with a 5 mm room-temperature BBO BB-H liquid NMR probe.

The sample temperature was controlled by a BCU-05 variable temperature unit. Temperature calibration was carried out with 4% MeOH in CD_3_OD. The stddiffesgp.3 pulse program from the Bruker pulseprogram library was used for data acquisition. The suppression of the HDO signal (4.703 ppm) was achieved by the excitation sculping method. The saturation frequency for the on- and off-resonances was set to −400 Hz and −16.000 Hz, respectively. The saturation time was 3 s with a relaxation delay D1 of 4 s.

The protein stock solutions were transferred to a deuterated buffer containing 35 mM potassium phosphate and 25 mM NaCl in D_2_O at a pD of 7.5 using 0.5 mL Centricon devices (Merck Millipore) with a MWCO of 30 kDa by washing and centrifuging several times at 10,000 x g and 4°C. Buffer-exchanged protein was adjusted to a concentration of 30 µM with NMR buffer. The respective fragment concentration was 428 µM with a final DMSO-d_6_ fraction of 5% (v/v). All measurements were performed in the presence of 500 µM ADP and 5 mM MgCl_2_. All data were analyzed using TopSpin 4.0.8 (Bruker Corporation, Billerica, MA, USA).

### HSQC NMR measurements

^15^N-labelled p97-N was concentrated to 50-60 µM in 25 mM HEPES (pH 7.5), 125 mM NaCl, 5 mM DTT and 0.01% NaN_3_. Before the measurements were conducted, 10% D_2_O and 3-(trimethylsilyl)propane-1-sulfonate (DSS) at a final concentration of 0.1 mM were added to the samples. DSS was used for direct ^1^H chemical shift referencing as 0.00 ppm. ^15^N chemical shifts were indirectly referenced by the gyromagnetic ratio [69].

^1^H-^15^N-HSQC spectra were recorded on a Bruker Avance III (600 MHz) spectrometer at 298 K. DMSO-dissolved ligands were added to the proteins, resulting in molar ratios of 1:80 or 1:160 (protein:ligand). As a control, equivalent volumes of pure DMSO were added. Assignments were obtained from previously recorded spectra of the p97 N domain [70,71]. Spectra were plotted and analysed using CcpNMR Analysis [72,73] version 3.0.4. Chemical Shift Perturbations (CSP) were calculated according to the equation:

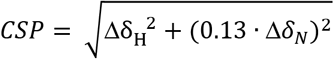

### Data availability

Atomic coordinates and structure factors have been deposited in the Protein Data Bank with accession code 7PUX.

## Acknowledgements and Funding

This work was supported by the Deutsche Forschungsgemeinschaft (GRK2243 and EXC 2051 - Project ID 390713860) and by the Rudolf Virchow Center for Integrative and Translational Bioimaging (P.H. and H.S.). The fragment screen was performed at the facility for Biophysics, Structural Biology and Screening at the University of Bergen (BiSS), which received funding from the Research Council of Norway (RCN) through the NOR–OPENSCREEN consortium (grant number 245922). We thank Khanh Dim Dao and Emil Hausvik for assistance with the screening; Markus Zehe and Curd Schollmayer (Institute of Pharmacy and Food Chemistry) for assistance with NMR measurements, Monika Kuhn (Rudolf Virchow Center) for assistance in protein expression and purification, and the beamline scientists at the ESRF. We thank the Centre of Biomolecular Magnetic Resonance (BMRZ) at the Goethe University Frankfurt funded by the state of Hesse for support.

## Author contributions

C.S., H.S. and P.H. initiated and supervised the project. S.B., P.H. and St.B. performed cloning, protein expression and purification. R.B. initiated fragment screening. S.B. performed BLI fragment screening and data analysis. S.B. and J.K. performed MD simulations and analyses. S.B. performed STD-NMR and CSP analysis and crystallization. S.B. and H.S. performed data collection and X-ray data analysis. St.B., U.A.H. and C.W. performed HSQC-NMR measurements. S.B. drafted the manuscript and all authors contributed additions and corrections. All authors have read and approved the final manuscript.

