## Supplemental material for "Fragment screening using biolayer interferometry reveals ligands targeting the SHP-motif binding site of the AAA+ ATPase p97"

### Supplementary Figures

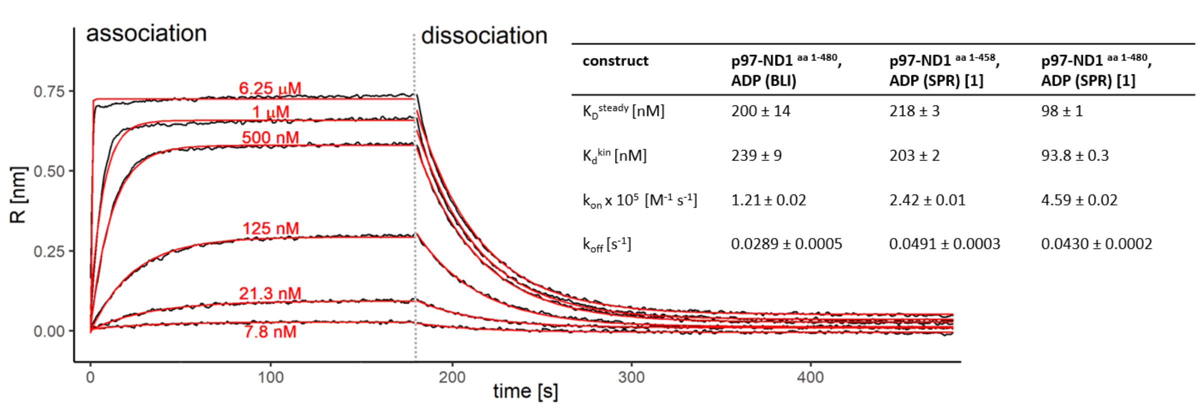

| construct | p97-ND1 aa 1-480, ADP (BLI) | p97-ND1 aa 1-458, ADP (SPR) [1] | p97-ND1 aa 1-480, ADP (SPR) [1] |
|---|---|---|---|
| $K_D^{steady}$ [nM] | 200 ± 14 | 218 ± 3 | 98 ± 1 |
| $K_d^{kin}$ [nM] | 239 ± 9 | 203 ± 2 | 93.8 ± 0.3 |
| $k_{on}$ x $10^5$ [$M^{-1}$ $s^{-1}$] | 1.21 ± 0.02 | 2.42 ± 0.01 | 4.59 ± 0.02 |
| $k_{off}$ [$s^{-1}$] | 0.0289 ± 0.0005 | 0.0491 ± 0.0003 | 0.0430 ± 0.0002 |

**Supplementary Figure 1. Validation of BLI assay with p97-ND1.** Shown are the sensorgrams and the global fitting model of ADP as positive control for assay validation. The sensorgrams clearly follow a 1:1 global binding model. For comparison, the affinity and kinetic constants of ADP for two similar p97-ND1 constructs determined via SPR by Chou et al. [1] are listed in the table insert.

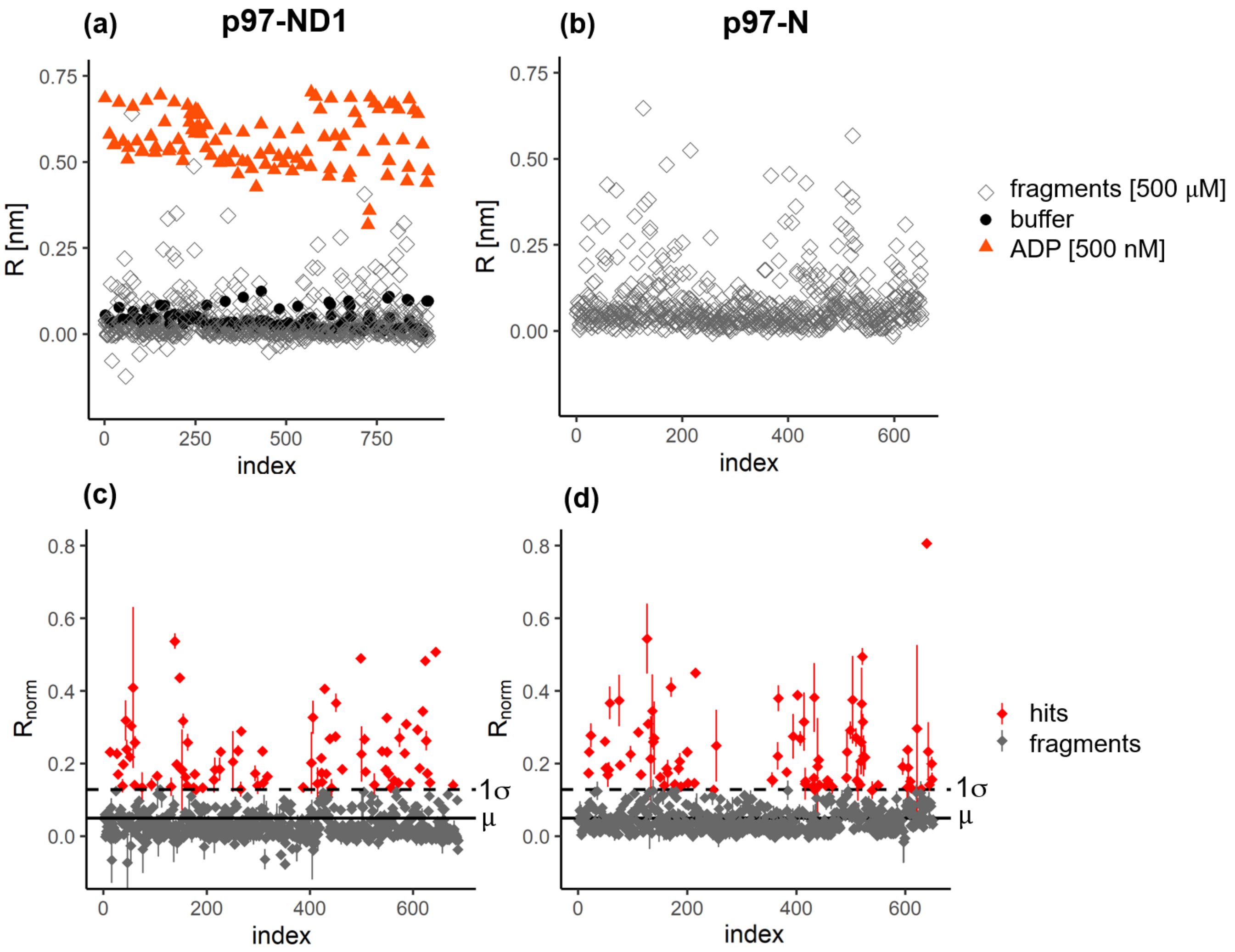

**Supplementary Figure 2. Initial BLI screen with the ND1 domain (left panels) and the N-domain (right panels) of p97. (a)** Results of the ND1 screen after double referencing. The signals of the ADP positive controls (red) and of the negative controls using assay buffer (black) are shown. Gray squares indicate the mean signal (n=2) of each measured fragment at a concentration of 500 µM. The numbers on the x-axis count fragments and positive controls and are therefore higher than in the subsequent diagrams with fragments only. **(b)** Results of the initial screen with the N-domain after double referencing and elimination of artifacts showing negative or extremely high values. Gray squares represent the mean signal (n=2) of each measured fragment at a concentration of 500 µM. **(c)** Normalized signals of the fragments of the ND1-domain screen with respect to the positive control, resulting in a mean normalized signal of 0.050 and a standard deviation of 0.079. **(d)** Normalized signals of the fragments of the N-domain screen with respect to the protein loading signals of the used sensors, giving a mean value of 0.069 and a standard deviation of 0.080. A cut-off of one sigma was used in both cases (dashed lines).

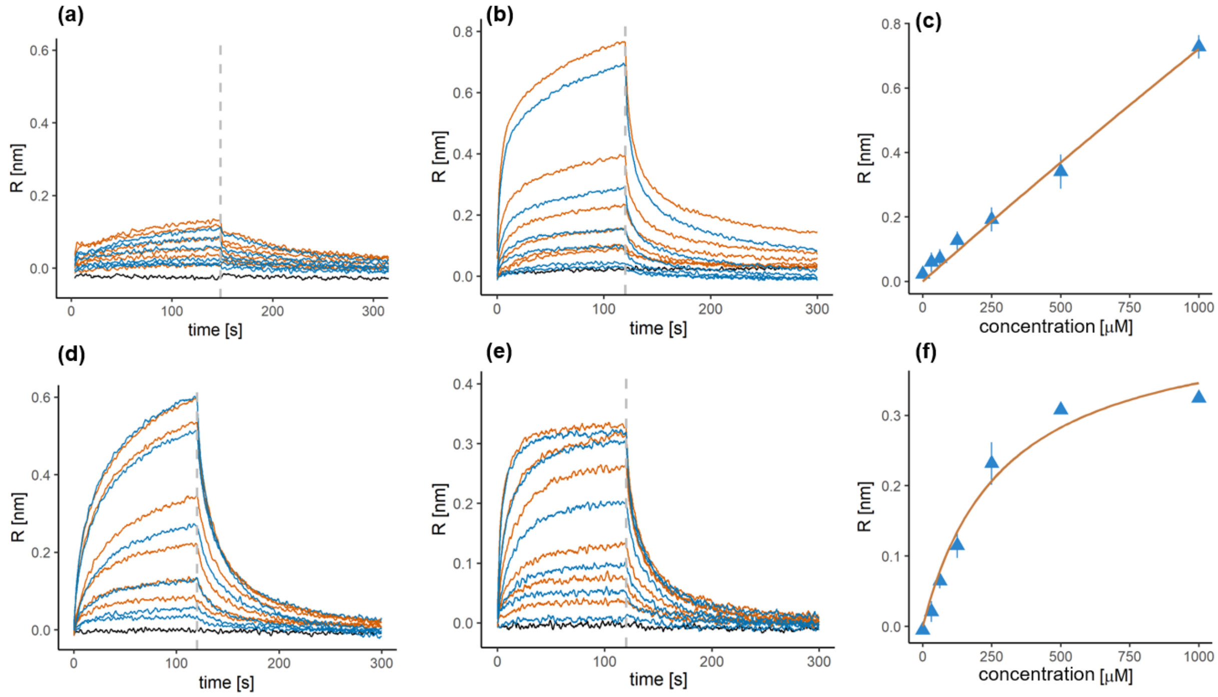

**Supplementary Figure 3. Representative binding profiles for the interaction of four compounds with p97-N.** Sensorgrams obtained with BLI from a 1:1 dilution series of 6 steps starting at a concentration of 1 mM. All concentrations were measured twice (first: orange, second: blue). The dashed line indicates the start of the dissociation phase. **(a)** Example of a compound exhibiting no significant signal. **(b)** Example of a compound showing a non-saturating response with increasing concentrations, which results **(c)** in a typical linear increase of the signal. In these cases, complete dissociation could not be reached. **(d)** Example of a fragment exhibiting a biphasic curve with no plateau phase at higher concentrations. **(e)** Sensorgram of one of the best binders. The curves reach a steady-state and show the typical shark fin profile. The analysis with a 1:1 Langmuir model **(f)** reveals the expected hyperbola characteristic of a single binding site exhibiting saturation at high concentrations.

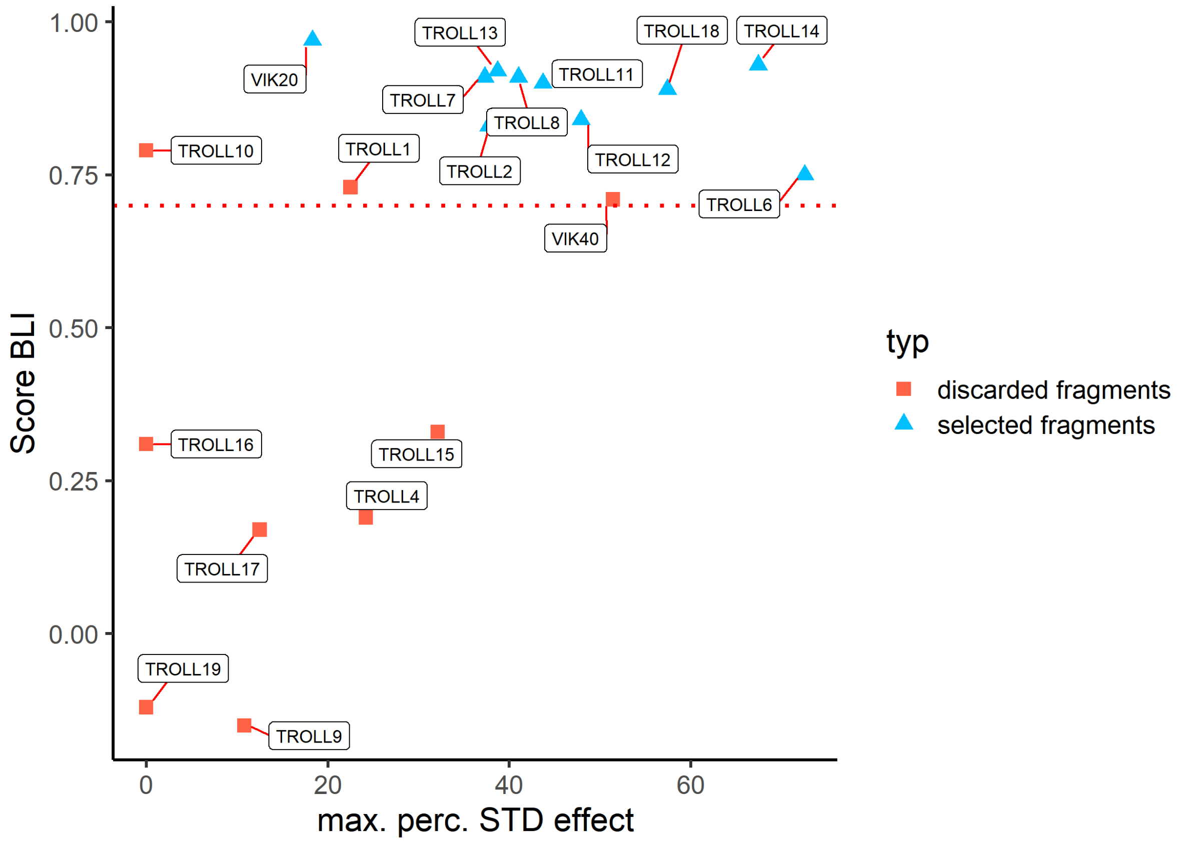

**Supplementary Figure 4. Overview of calculated Score$_{BLI}$ and measured STD-effects of the fragments.** Selected fragments are shown in blue and discarded molecules in red. Three fragments showed no STD-effect (TROLL10, TROLL16 and TROLL19). All other fragments displayed STD-effects between 10.5 and 72.5%. Selected fragments fulfill the criterion of a Score$_{BLI}$ value greater than 0.7. TROLL1 was discarded because no dose response with the p97 ND1-construct was obtained, VIK40 was discarded because of its low affinity ($K_D$ > 1 mM) with the ND1-construct and TROLL10 because of its high discrepancy in affinity determined either by the kinetic analysis or in steady state measurements.

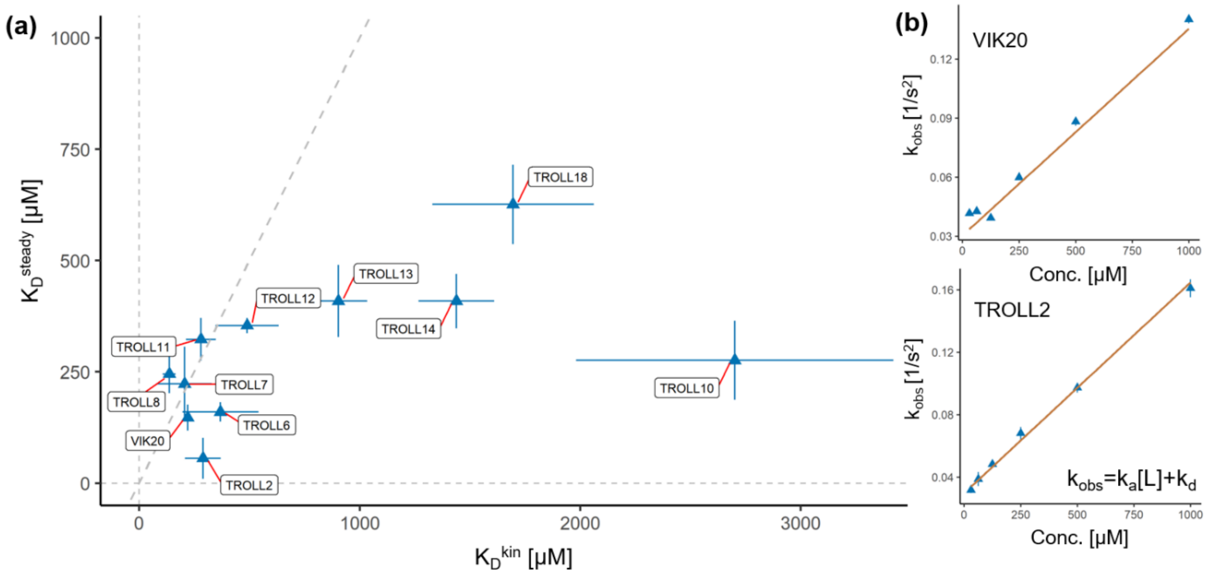

**Supplementary Figure 5. Comparison of kinetic and steady state determined affinity of the fragments.** Since the sensorgrams of the fragments showed slow on and off rates, the affinities were calculated based on the kinetic data of the dilution measurements. **(a)** Comparison of the kinetic affinity with the affinity determined by steady state. The diagonal shows the position of equal affinity values. Most fragments showed similar affinities. TROLL13, TROLL14 and TROLL18 showed a discrepancy of an up to 3.5 times lower affinity based on the kinetic data. The highest discrepancy was found for TROLL10 resulting in a 10-fold lower affinity derived from the kinetic data. **(b)** The on and off rate constants were determined using the empirical rate constants and their dependency on the used concentration under the assumption of a 1:1 binding. Plotted are the results for TROLL2 and VIK20, which show a good agreement with the 1:1 binding model.

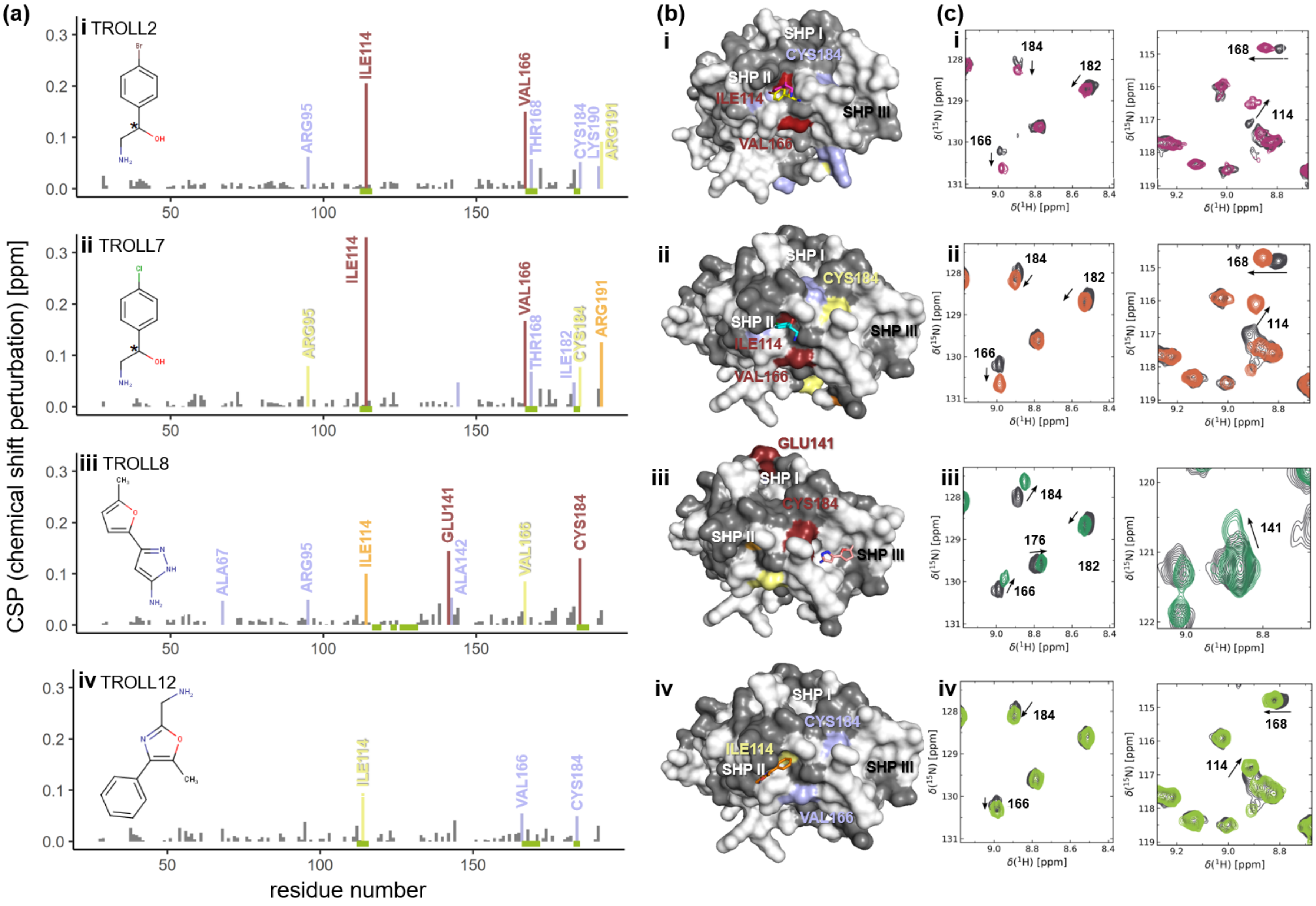

(a)
i TROLL2
ii TROLL7
iii TROLL8
iv TROLL12
CSP (chemical shift perturbation) [ppm]
residue number
ILE114
VAL166
THR168
ARG95
CYS184
LYS190
ARG191
ILE182
ALA67
GLU141
ALA142
(b)
SHP I
SHP II
SHP III
(c)
δ(¹H) [ppm]
δ(¹⁵N) [ppm]

**Supplementary Figure 6. Chemical shift perturbation (CSP) observed upon titration of the p97 N-domain with the corresponding fragment. (a)** Changes in the amide chemical shift are plotted for the p97 N-domain residues in terms of their strength corresponding to σ-levels of 4 (red), 3 (orange), 2 (yellow), 1 (blue). **(b)** Mapping of the chemical shifts onto the surface of the N-domain together with the binding poses of the fragments (stick representation) received from the mixed-solvent MD simulations. **(c)** Selected regions of the $^{1}H$-$^{15}N$-HSQC NMR spectra**.** Fragments in DMSO were titrated to $^{15}N$-labeled p97-N (colored spectra)**.** The respective residue numbers are indicated based on the established backbone assignment [1]. The control spectra upon addition of an equal amount of DMSO are shown in gray.

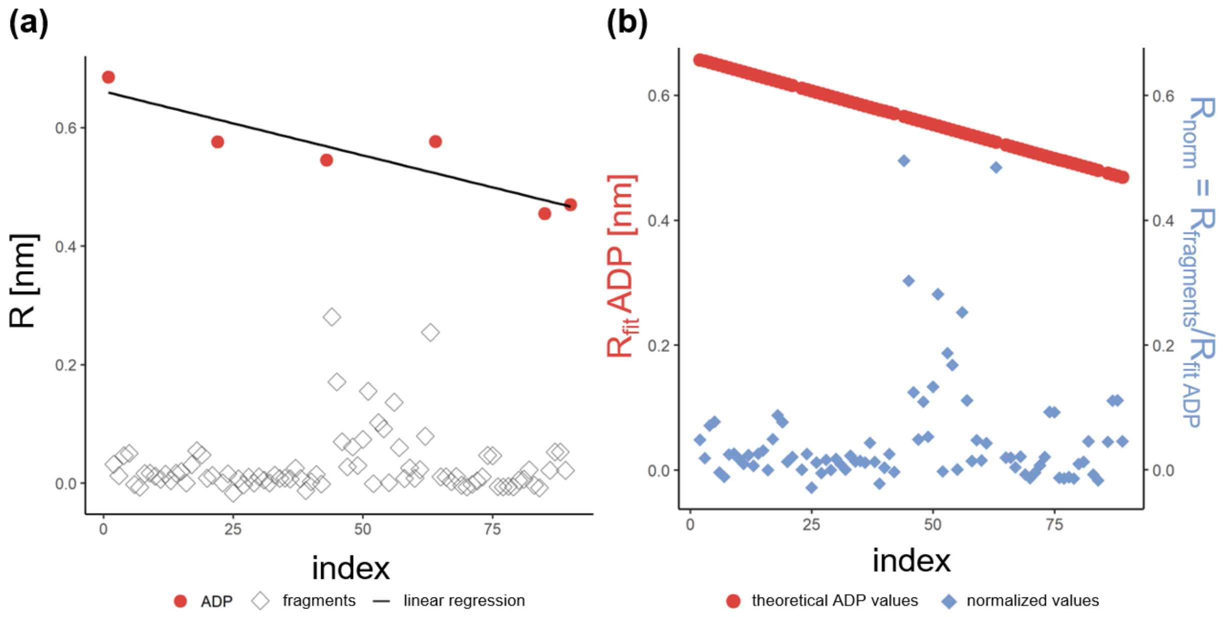

**Supplementary Figure 7. Normalisation of signals for the ND1 screen.** The signals of each sensor were corrected for partial inactivation of the protein over time and for different protein loading states of the sensors using 500 nM ADP as the positive control. Prior to normalization, the signals were aligned and double referenced using the Octet Analysis software. The normalization procedure is shown for one sensor: **(a)** With the signals of the ADP positive control (red dots) measured between each screening plate a linear regression was performed (black line). Grey diamonds reflect the signals of the fragments. **(b)** Using the obtained linear equation, the theoretically expected ADP signal at the time of measurement of each fragment was calculated (red dotted line). The normalization of the fragment signals was then carried out via $R_{norm}=R_{fragment}/R_{fit\ ADP}$ resulting in the normalized fragment values (blue diamonds).

### Supplementary Tables

**Table S1. Data collection and refinement statistics for p97-ND1 in complex with TROLL2** (PDB entry: 7PUX).

| **Data collection** | |
|---|---|
| Wavelength (Å) | 0.9184 |
| Resolution (Å) | 47.87-1.73 (1.83-1.73) [a] |
| Space group | P622 |
| Cell dimensions | |
| *a=b, c* (Å) | 146.23, 84.40 |
| *α=β, γ* (°) | 90, 120 |
| Unique reflections | 42,850 |
| <I/σ(I)> | 16.68 (1.63) [a] |
| Multiplicity | 26.47 |
| Completeness *spherical* (%) | 76.9 (25.31) [a] |
| Completeness *ellipsoidal* (%) | 96.12 (82.0) [a] |
| CC(1/2) | 0.999 (0.620)[a] |
| $R_{pim}$ | 0.027 (0.492) [a] |
| $R_{meas}$ | 0.140 (2.547) [a] |
| **Refinement statistics** | |
| Resolution limits | 47.87-1.73 |
| $R_{work}/R_{free}$ | 0.192/0.235 |
| Average B-factor (Å$^2$) | 37.3 |
| Root mean square deviations | |
| *Bond lengths* (Å) | 0.006 |
| *Bond angles* (°) | 0.89 |
| Ramachandran statistics (%) | |
| *Favored* | 97.5 |
| *Allowed* | 2.5 |
| *Outliers* | 0.0 |

[a] Numbers in parentheses refer to the highest resolution data shell

**Table S2. Comparison of the fragments with confirmed dose response from the screenings with the ND1-construct and the N-domain, respectively.** In addition, the mean and maximum STD-effect obtained in the NMR experiment with the ND1-construct is reported for selected compounds.

***n.b.*** no binding signal observed; ***n.d.*** affinity not determinable due to weak signal or poor signal quality; “-” not measured

| ID | structure | $K_D$ [µM] (ND1) | $K_D$ [µM] (N) | mean / maximum STD-effect (ND1) |
|---|---|---|---|---|
| TROLL1 | 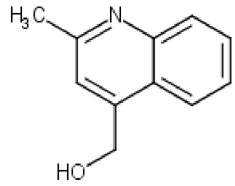 | n.b. | 37±3 | 18.0 / 22.5 |
| TROLL2 | 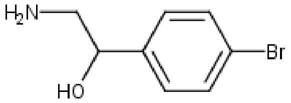 | 321±67 | 56±46 | 36.0 / 37.7 |
| VIK40 | 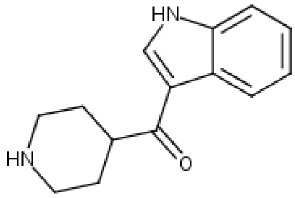 | 707±32 | 93±4 | 43.6 / 51.4 |
| TROLL4 | 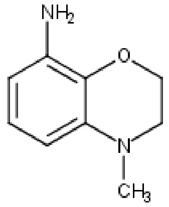 | 292±135 | 124±29 | 20.5 / 24.2 |
| VIK20 | 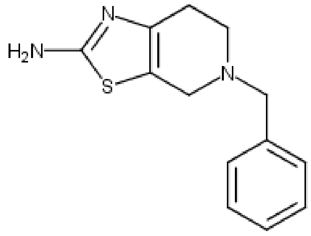 | 192±92 | 147±32 | 18.3 / 18.3 |
| TROLL6 | 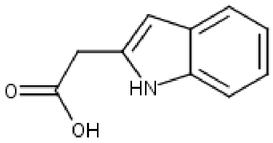 | 243±16 | 160±22 | 66.5 / 72.5 |
| TROLL7 | 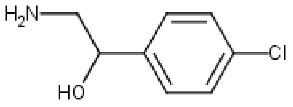 | 718±57 | 222±83 | 34.5 / 37.3 |
| TROLL8 | 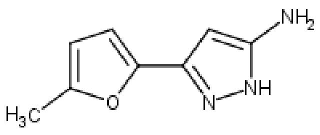 | 598±214 | 245±43 | 33.1 / 41.0 |

| ID | structure | $K_D$ [μM] (ND1) | $K_D$ [μM] (N) | mean / maximum STD-effect (ND1) |
|---|---|---|---|---|
| TROLL9 | 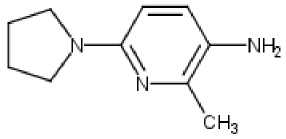 | 351±82 | 267±65 | 10.5 / 10.8 |
| TROLL10 | 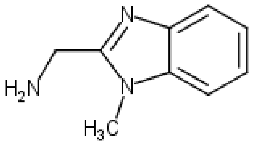 | 66±14 | 276±89 | n.b. |
| TROLL11 | 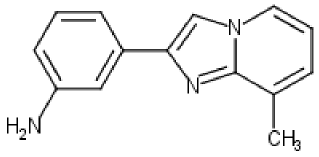 | 290±179 | 323±48 | 37.8 / 43.7 |
| TROLL12 | 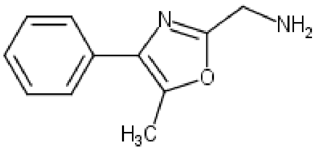 | 192±92 | 354±17 | 44.5 / 47.9 |
| TROLL14 | 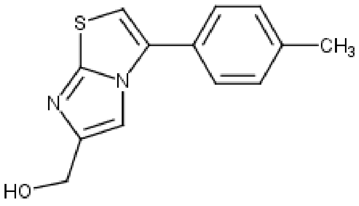 | 290±33 | 409±61 | 55.4 / 67.4 |
| TROLL13 | 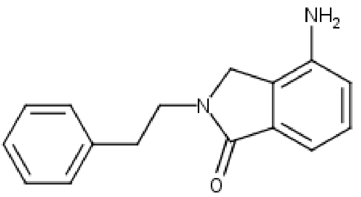 | 784±118 | 409±80 | 33.0 / 38.7 |
| TROLL15 | 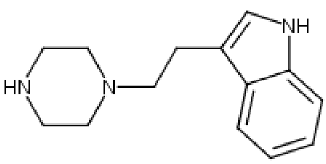 | 303±82 | 425±31 | 25.2 / 32.1 |
| TROLL16 | 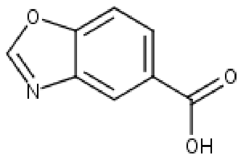 | n.d. | 440±53 | n.b. |
| TROLL17 | 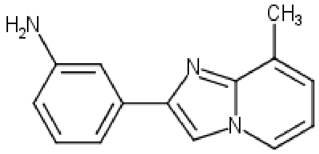 | n.b. | 444±56 | 11.3 / 12.5 |
| TROLL18 | 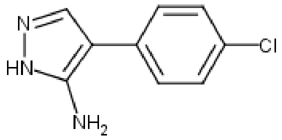 | 445±223 | 625±89 | 54.3 / 57.4 |

| ID | structure | $K_D$ [µM] (ND1) | $K_D$ [µM] (N) | mean / maximum STD-effect (ND1) |
|---|---|---|---|---|
| TROLL19 | 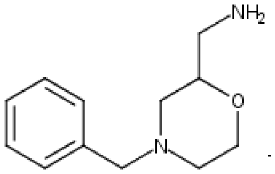 | 541±25 | 641±125 | n.b. |
| TROLL20 | 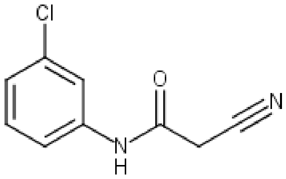 | n.d. | 714±525 | - |
| TROLL21 | 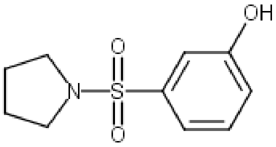 | 62±31 | 821±101 | - |
| VIK50 | 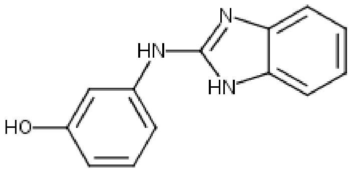 | 192±92 | 853±57 | 73.9 |
| VIK80 | 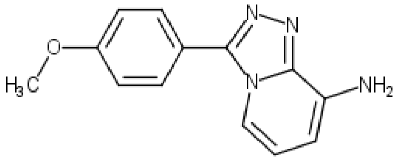 | 112±16 | - | - |
| VIK1 | 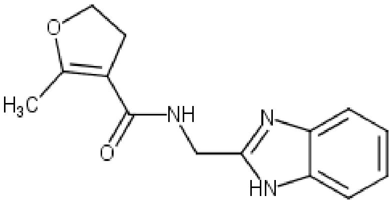 | 115±13 | - | - |
| ODIN27 | 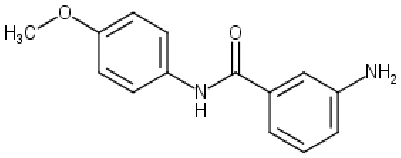 | 143±5 | - | - |
| ODIN32 | 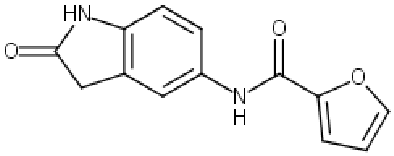 | 178±20 | - | - |
| ODIN33 | 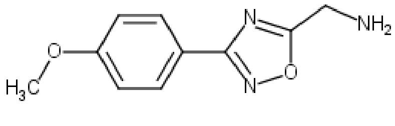 | 281±19 | - | - |

| ID | structure | $K_D$ [µM] (ND1) | $K_D$ [µM] (N) | mean / maximum STD-effect (ND1) |
|---|---|---|---|---|
| ODIN26 | 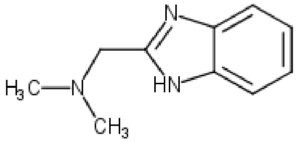 | 290±170 | - | - |
| ODIN10 | 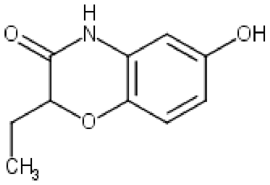 | 367±58 | - | - |
| VIK110 | 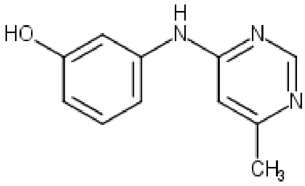 | 428±33 | - | - |
| ODIN18 | 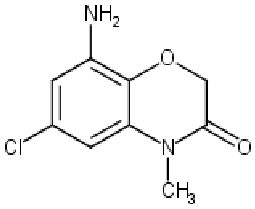 | 657±89 | - | - |
| ODIN13 | 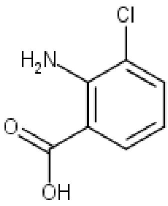 | 723±57 | - | - |
| VIK90 | 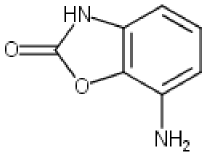 | 963±408 | - | - |

*Sorted by $K_D$ (N), and subsequently by $K_D$ (ND1).*

### Supplementary Methods

#### Definition of the quality score "Score$_{BLI}$"

For selecting fragments for further analysis, the $Score_{BLI}$ was defined as:

$$Score_{BLI} = \frac{r^2_{mean\ 1:1}}{r^2_{mean\ 2:1}}\ r^2_{kobs} - 100|m_{Plateau}|\ (1)$$

$r^2_{mean\ 1:1}$ is the mean of the $r^2$ coefficients of the regressions of all association phases measured for a given fragment. The data were fitted with a 1:1 binding model:

$$R_t = R_{eq}\left(1 - e^{-(kobs\ t)}\right)\ (2)$$

with $R_t$ the shift at the time *t*, $k_{obs}$ the empirical rate constant and $R_{eq}$ the shift at equilibrium. In analogy, the $r^2_{mean\ 2:1}$ was defined by fitting with a 2:1 binding model:

$$R_t = R_{eq\ 1}\left(1 - e^{-(kobs1\ t)}\right) + R_{eq\ 2}\left(1 - e^{-(kobs2\ t)}\right)\ (3)$$

A ligand following a 1:1 model would result in a good $r^2_{mean}$ for both functions, whereas for a ligand which is better described with a 2:1 model the $r^2_{mean}$ for a 1:1 model would decrease, resulting in an overall decreased value.

In case of a 1:1 model the empirical rate constant $k_{obs}$ is proportional to the concentration of the ligand *[L]*:

$$k_{obs} = k_a[L] + k_d\ (4)$$

with $k_a$ the association rate constant and $k_d$ the dissociation rate constant. Therefore, all observed empirical rate constants of a ligand were subjected to a linear regression model, which yielded the value $r^2_{kobs}$.

For the estimation of the affinities of the fragments, a Langmuir model was used. The use of this model requires that a chemical equilibrium is reached. To analyze whether this condition was met, the responses of the last 10 seconds of each association phase were fitted with a linear model. In the chemical equilibrium the slope should be zero. The obtained slopes of each concentration were plotted against the logarithm of the molar concentration and fitted with a linear model. If the equilibrium was successfully reached for all concentrations, the resulting slope $m_{plateau}$ should also be close to zero, whereas unspecific binding, where no chemical equilibration can be observed, will lead to a slope > zero. The resulting value $m_{plateau}$ was used as a "penalty term". The calculated $Score_{BLI}$ should be close to or above 1.0 for a good 1:1 ligand, whereas ligands showing unspecific binding should yield values < 1.0. The score was evaluated with binding data for ADP, which gave a $Score_{BLI}$ of 1.02, and for a clearly non-specifically binding fragment (TROLL9), which gave a $Score_{BLI}$ of -0.15.
